# Podocyte-to-Parietal Epithelial Cell Senescence Drives Experimental Focal Segmental Glomerulosclerosis and is reversed by Dasatinib Plus Quercetin

**DOI:** 10.64898/2026.11.26.769476

**Authors:** Antonio Battaglia-Vieni, Cristina Garcia-Caballero, Fátima Milhano Santos, Tri.Q. Nguyen, Nerea Mendez-Barbero, Sergio Ciordia, Alberto Ortiz, Anna Sola, Sandra Rayego-Mateos, Marta Ruiz-Ortega

**Affiliations:** Molecular and Cellular Biology in Renal and Vascular Pathology. Department of Medicine, Universidad Autónoma de Madrid, IIS-Fundación Jiménez Díaz, Madrid, Spain; Ricors2040. Instituto de Salud Carlos III, Madrid Spain; Department of Pathology, UMC Utrecht Netherlands; Vascular Research Laboratory, IIS-Fundación Jiménez Díaz, CIBERCV, ISCIII, Madrid, Spain; Faculty of Biomedical Science and Health, Universidad Alfonso X el Sabio (UAX), Madrid, Spain; Laboratory of Functional Proteomics, Centro Nacional de Biotecnología, CSIC, Madrid, Spain; Department of Nephroloy, IIS-Fundación Jiménez Díaz, Universidad Autónoma de Madrid, Madrid Spain; Department of Nephrology and Transplantation, Institut d’Investigació Biomèdica de Bellvitge (IDIBELL), Hospitalet de Llobregat, Barcelona, Spain; GLOMED (Glomerular medicine and therapeutics group); IIS-Fundación Jiménez Díaz, Madrid, Spain

**Keywords:** Focal segmental glomerulosclerosis, Cellular senescence, Podocyte, Parietal epithelial cell, Senolytics, Dasatinib, Quercetin

## Abstract

Focal segmental glomerulosclerosis (FSGS) is a progressive podocyte-depleting glomerular disease and a leading cause of chronic kidney disease, yet the contribution of cellular senescence to its pathogenesis remains poorly understood. Here, we identified senescent parietal epithelial cells (PECs) in human FSGS, revealing a previously unrecognized component of glomerular injury. In experimental FSGS, podocyte senescence preceded PEC senescence, which was associated with glomerulosclerosis. Conditioned medium from senescent podocytes induced secondary senescence in PECs. In preclinical FSGS, therapeutic administration of senolytics dasatinib plus quercetin (DQ) reduced senescent cells and activation of related signaling pathways, attenuated glomerular injury, and improved kidney function. Proteomic profiling identified activation of interferon and STAT3 pathways in FSGS kidneys modulated by DQ treatment. These findings identify podocyte-to-PEC senescence signaling as a novel mechanism promoting glomerular injury and sclerosis, support cellular senescence as a therapeutically modulable pathway, and provide a translational rationale for testing senolytic therapies in FSGS.

## Introduction

Focal segmental glomerulosclerosis (FSGS) is a podocyte-depleting pattern of glomerular injury and a leading cause of nephrotic syndrome and chronic kidney disease (CKD). Despite advances in diagnosis and treatment, FSGS remains a major clinical challenge because of its heterogeneous clinical course and highly variable therapeutic responses ^1–4^. Therefore, there is an unmet clinical need to identify mechanisms underlying glomerular injury and disease progression and to develop targeted interventions that improve disease outcomes.

Podocyte injury is the hallmark of FSGS^3^. However, the mechanisms linking podocyte loss to progressive glomerular scarring remain poorly understood. Podocyte depletion disrupts the integrity of the glomerular filtration barrier, promoting proteinuria and pathological repair responses that accelerate sclerosis and kidney function decline^3^. Recent evidence has challenged the traditional view of parietal epithelial cells (PECs) as passive structural components of Bowman’s capsule, highlighting their role as important regulators of glomerular homeostasis and contributors to glomerular injury and disease progression^5^. In FSGS, the initial podocyte injury is followed by PEC activation and phenotypic reprogramming, characterized by de novo CD9 and CD44 expression, enhanced proliferative and migratory capacity, and increased extracellular matrix production. Through these maladaptive responses, PECs have emerged as major drivers of FSGS progression ^6–9^. Recent studies highlight cell-to-cell communication as an important pathogenic mechanism in FSGS, as injured podocytes release paracrine signals that induce PEC activation and promote glomerular scarring^10^. These observations suggest that pathogenic podocyte-to-PEC crosstalk constitutes a previously unrecognized mechanistic link between podocyte loss and progressive glomerulosclerosis. However, the molecular mediators governing this process remain largely unknown.

Cellular senescence has emerged as a fundamental mechanism linking tissue injury to chronic inflammation and fibrosis^11^. Senescent cells undergo irreversible cell-cycle arrest and upregulate anti-apoptotic pathways, while remaining metabolically active and secreting a complex pro-inflammatory and pro-fibrotic senescence-associated secretory phenotype (SASP) that can induce cellular dysfunction and secondary senescence in neighboring cells^12–14^. Senescence may arise as a primary response to cell-intrinsic stress or develop as secondary senescence through the paracrine and juxtacrine actions of SASP components^15^. Senescent cell accumulation and persistent SASP signaling are now recognized as key drivers of CKD progression ^16^. Importantly, senescence can be therapeutically targeted through senolytic strategies that selectively eliminate senescent cells, attenuating kidney injury and fibrosis^17^. Podocyte senescence has been implicated in the pathogenesis of glomerular diseases by limiting podocyte regenerative capacity and promoting local inflammation through SASP-mediated signaling^18^. Nevertheless, whether senescence-dependent propagation of pathogenic signals from podocytes to PECs contributes to glomerulosclerosis and FSGS progression remains unknown.

Among the available senolytic strategies, the combination of dasatinib plus quercetin (DQ) has emerged as one of the most extensively studied approaches. DQ has demonstrated beneficial effects in ageing-related disorders^19^ and experimental models of CKD, including diabetic kidney disease^19–22^. However, DQ efficacy appears to be highly context-dependent, varying with disease type, timing of intervention, and underlying pathogenic mechanisms^23,24^ highlighting the need for disease-specific evaluation in kidney disorders.

Given the lack of effective disease-modifying therapies for FSGS and the emerging role of cellular senescence in kidney diseases, we hypothesized that senescence propagates within the glomerular niche through paracrine communication between podocytes and PECs, thereby amplifying glomerular injury and promoting FSGS progression. To address this hypothesis, we characterized glomerular senescence in human and experimental FSGS, investigated the role of senescence-associated podocyte-PEC communication, and assessed the therapeutic potential of senolytic intervention with DQ.

## Methods

Detailed information is available in Supplementary Methods.

### Ethical statement

Experiments on human samples followed all relevant guidelines and regulations. The use of kidney biopsies for research purposes was approved by the Biobank Research Ethics Committee of UMC Utrecht, Netherlands (TCBio 19-945 269). Patients provided written informed consent. The studies were conducted in accordance with the local legislation and institutional requirements and the Declaration of Helsinki.

Animal procedures were performed in mice according to the European Community and ARRIVE reporting guidelines for the care and use of laboratory animals, after approval by the Animal Ethics Committee of the IIS-Fundación Jiménez Díaz and Comunidad Autónoma de Madrid, Spain Madrid (PROEX 0019.0/26).

### Human samples

Six human kidney biopsies with FSGS were selected from the pathology archives of UMC Utrecht and immunohistochemistry for p21Cip1 was performed using a Ventana RTU antibody DCS-60.2 (Cell Marque, Rocklin, CA, USA).

### Animal models

BALB/c mice (male except indicated) received a single retro-orbital injection of adriamycin (ADR; 10 mg/kg) to induce FSGS ^25^. Senolytic combination DQ (dasatinib 5 mg/kg plus quercetin 50 mg/kg) was administered via oral gavage starting on day 3 post-injury and continued every 4 days.

## Results

### Human FSGS is characterized by podocyte and PEC senescence

Because cellular senescence is a heterogeneous and context-dependent state, with substantial variability in the expression of individual senescence markers across cell types and pathological conditions, we first sought to determine whether human FSGS exhibits a broader senescence-associated molecular signature. We therefore assessed the expression of canonical senescence markers and components of the SASP in publicly available human FSGS transcriptomic datasets using Nephroseq v5. Expression of the cell-cycle inhibitors p21^Cip1^ (*CDKN1A)* and p16^INK4a^ (*CDKN2A*) correlated with renal function decline in FSGS (Supplementary figure 1A), while multiple SASP-associated genes were significantly upregulated in glomeruli from patients with FSGS (Supplementary figure 1B and 1C), revealing a senescence-associated molecular signature in human FSGS.

Next, we determined the cellular localization of this senescence-associated response in human FSGS kidney biopsies. We focused on p21^Cip1^, a key mediator of senescence-associated cell-cycle arrest and a well-established marker of cellular senescence in kidney disease and aging ^20,26^. Previous studies have demonstrated increased p21^Cip1^ expression in podocytes from patients with FSGS^18,27,28^. Consistent with these observations, p21^Cip1^-positive podocytes were detected in affected glomeruli from patients with FSGS (Figure 1a). Notably, we also identified p21^Cip1^-positive PECs within segmentally sclerosed glomeruli, whereas no p21^Cip1^-positive cells were observed in unaffected glomeruli (Figure 1b and 1c). These findings identify PECs as a previously unrecognized senescence-associated cell population within the sclerotic glomerular lesion.

**Figure 1.**
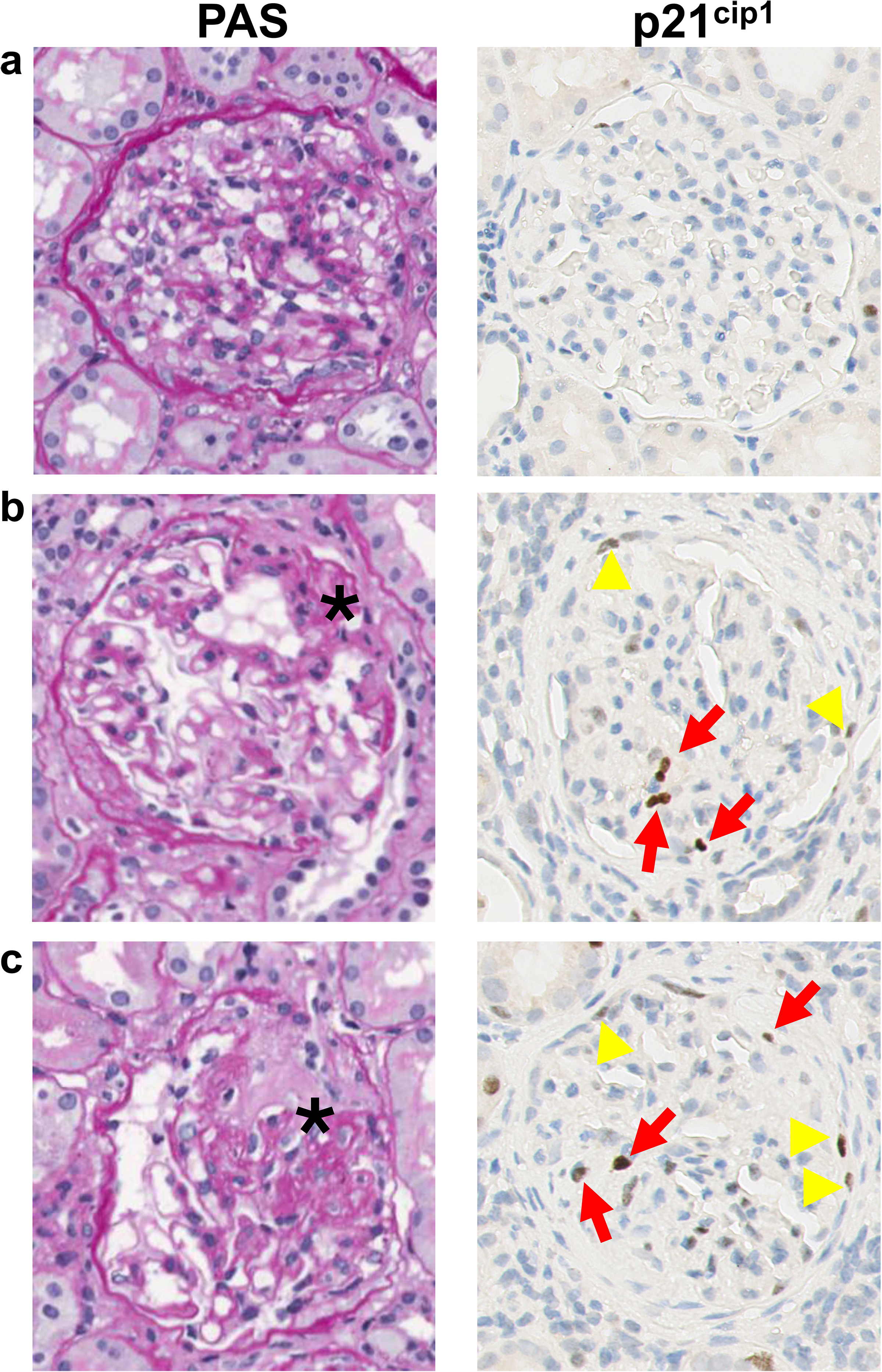
Senescent podocytes and parietal epithelial cells accumulate in human focal segmental glomerulosclerosis (FSGS). a, Representative image of an unaffected glomerulus from a 62-year-old individual with FSGS showing preserved glomerular architecture by PAS staining and no detectable p21^Cip1^-positive cells. b,c, Representative images from the same kidney biopsy showing affected glomeruli with segmental sclerosis (asterisks) and p21^Cip1^-positive cells in both podocyte and parietal epithelial cell compartments (closed and open arrows, respectively), indicating the presence of senescent PECs in human FSGS lesions.

### Podocyte senescence precedes PEC senescence during experimental FSGS progression

In ADR-induced experimental FSGS, podocyte senescence preceded the appearance of senescent PECs, suggesting a temporal hierarchy in glomerular senescence. On day 5 after ADR administration, p21^Cip1^-positive cells were rarely detected. By day 15, senescence was largely restricted to podocytes, whereas by day 22 it was evident in both podocyte and PEC compartments, coinciding with progressive glomerulosclerosis (Figure 2a and 2b). Co-staining with the PEC activation marker CD44^6^ confirmed the presence of senescent PECs at advanced stages of disease (Figure 2c). Quantitative analysis demonstrated a progressive increase in positive senescent cells over time (Figure 2d). In line with these findings, *Cdkn1a* gene expression was significantly increased in whole-kidney samples, as well as in isolated glomeruli from ADR-treated mice (Figure 2e and 2f).

**Figure 2.**
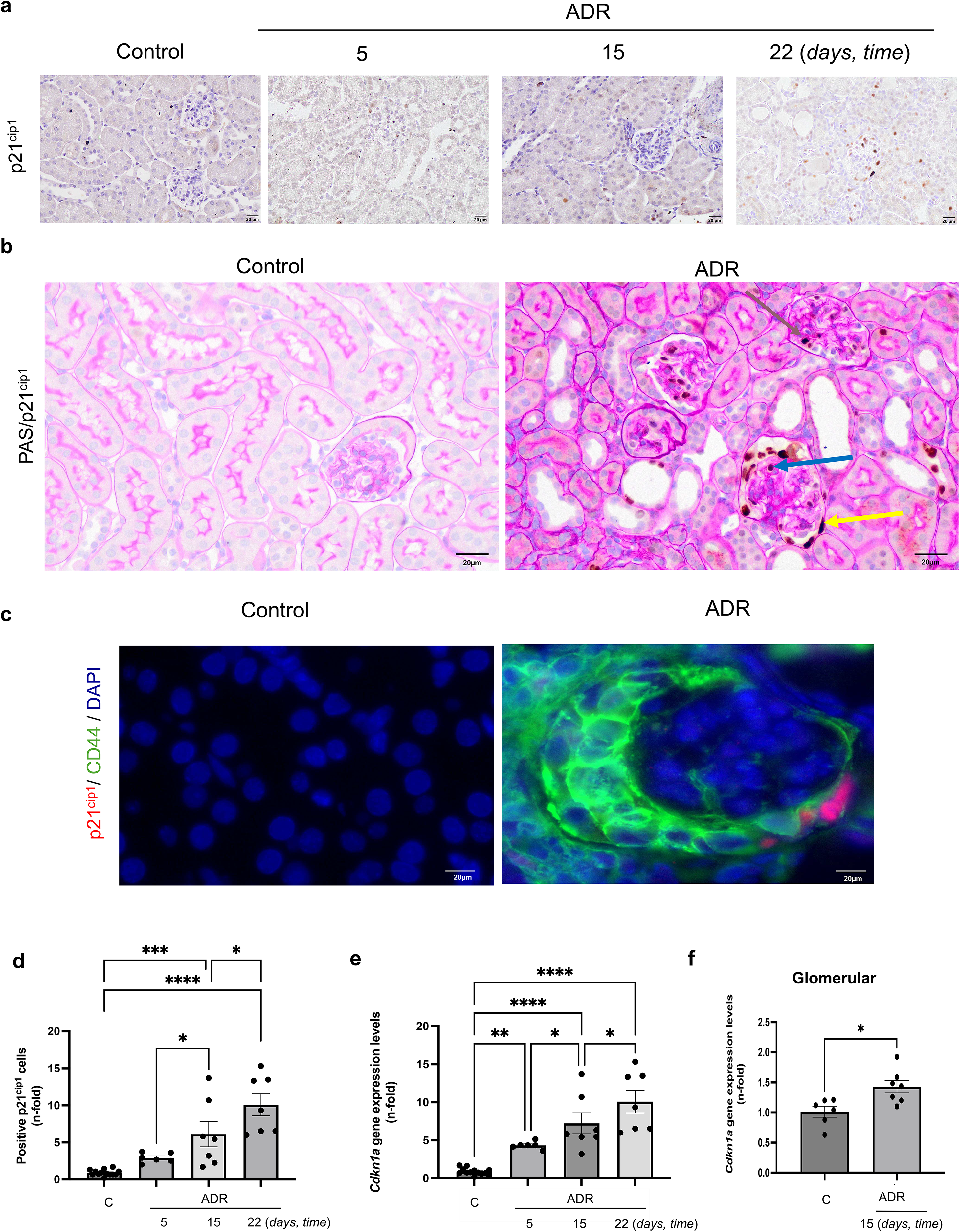
Glomerular senescence expands from podocytes to parietal epithelial cells (PECs) during experimental focal segmental glomerulosclerosis (FSGS) progression. Male BALB/c mice received a single retro-orbital injection of adriamycin (ADR; 10 mg/kg) to induce. a, Representative p21^Cip1^ immunohistochemistry in kidneys from control mice and ADR-treated mice at days 5, 15 and 22 (40× magnification). Arrows indicate p21^Cip1^-positive podocytes (blue) and p21-positive PECs (yellow). b. Representative PAS/p21^Cip1^ double staining showing the spatial distribution of senescent cells within the glomerulus during disease progression. Scale bars, 20 μm. c, Co-localization of p21^Cip1^ and the PEC activation marker CD44 identifies senescent PECs in advanced stages of experimental FSGS. Representative immunofluorescence images of p21^Cip1^ (red), CD44 (green) and DAPI (blue) in kidneys from 22 days ADR-treated mice. Scale bars, 100 μm. d, Ǫuantification of p21^Cip1^-positive cells at days 5, 15 and 22 after ADR administration, expressed as fold change relative to control mice. e,f, RT–qPCR analysis of *Cdkn1a* expression in (e) total renal cortex and (f) isolated glomeruli from control and ADR-treated mice, normalized to *Gapdh*. Data are presented as mean ± s.e.m; each symbol represents one mouse. n=6-13 control, n=6 ADR day 5, n=7 ADR day 15 and n=7 ADR day 22. p21^Cip1^-positive cells were analyzed using the Kruskal–Wallis test followed by Fisher’s LSD test. Renal cortex gene expression was analysed by ordinary one-way ANOVA with Fisher’s LSD post hoc test, whereas glomerular expression was analysed using a parametric unpaired two-tailed t-test. *p<0.05, **p<0.005, ***p<0.0005.

### Senescent podocytes induce secondary senescence in PECs

Exposure of podocytes to ADR induced a senescent phenotype, as demonstrated by increased senescence-associated β-galactosidase (SA-β-gal) activity in primary cultures, which was blocked by DQ treatment (Figure 3a and 3b). Accordingly, ADR-exposed human podocytes exhibited increased nuclear accumulation of p21^Cip1^ assessed by immunofluorescence (Figure 3c), together with elevated p21^Cip1^ protein and *CDKN1A* mRNA expression (Figure 3d and 3e), consistent with activation of a senescence-associated cell-cycle arrest program, as previously described^29^. These results showed that the treatment with DQ reduced the ADR-induced podocyte cellular senescence program, demonstrating its senolytic activity.

**Figure 3.**
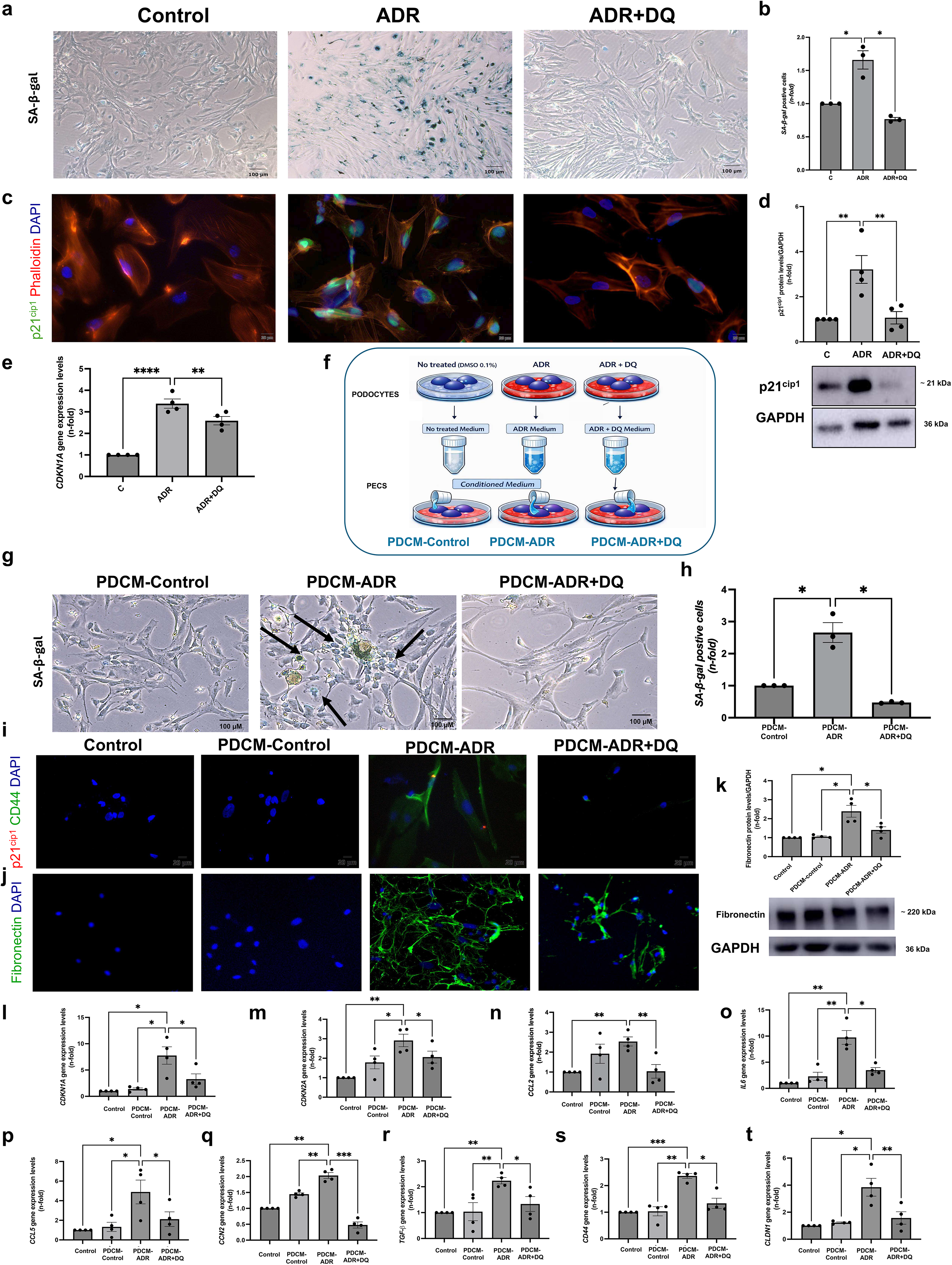
In vitro modelling reveals ADR-induced podocyte senescence as a driver of secondary senescence in PECs. a, Representative senescence-associated β-galactosidase (SA-β-gal) staining of primary murine podocytes following adriamycin treatment (ADR; 1 μg/mL, 24 h). A subset of cultures received dasatinib (600 nM) and quercetin (30 μM) (DǪ). b, Ǫuantification of SA-β-gal-positive podocytes, expressed as fold change relative to untreated controls. c, Representative p21^Cip1^ immunofluorescence in cultured human podocytes following ADR exposure. d, Immunoblot analysis of p21^Cip1^ protein expression in human podocytes, using GAPDH as loading control. e, RT–qPCR analysis of *CDKN1A* expression in human podocytes. f, <u>Experimental design of secondary senescence studies</u>. Podocytes were exposed to ADR (1 μg/mL) with or without dasatinib (600 nM) and quercetin (30 μM) (DǪ) for 24 h, followed by transfer of podocyte-derived conditioned medium (PDCM) to PEC cultures. g,h, Representative SA-β-gal staining (g) and quantification (h) of primary murine PECs exposed to PDCM. i, Representative immunofluorescence showing p21^Cip1^ and CD44 expression in PECs. Fibronectin expression assessed by immunofluorescence (j) and immunoblotting (k). l–t, RT–qPCR analysis of senescence-associated genes (*CDKN1A, CDKN2A, CCL-2, ILC, CCL5, CCN2* and *TGFB1*) and PEC markers (*CD44* and *CLDN1*) in human PECs exposed to control medium or PDCM. Data are presented as mean ± s.e.m. from 3–4 independent experiments. Statistical analyses were performed using one-way ANOVA followed by Fisher’s LSD post hoc test. *p<0.05, **p <0.005, ***p<0.0005.

To investigate whether senescent podocytes promote secondary senescence in PECs, podocyte-derived conditioned medium (PDCM) was collected from control podocytes, ADR-induced senescent podocytes, or ADR-treated podocytes receiving the senolytic combination DQ, and PDCM was subsequently applied to PEC cultures (Figure 3f). Exposure of primary murine PECs to PDCM from ADR-induced senescent murine podocytes significantly increased SA-β-gal activity, suggesting the induction of secondary senescence in these cells (Figure 3g and 3h). Accordingly, exposure to PDCM from senescent human podocytes induced a robust senescence program in human PECs, as evidenced by increased p21^Cip1^ and CD44 immunofluorescence (Figure 3i) and the upregulation at gene levels of the cell-cycle arrest markers such as *CDKN1A* and *CDKN2A* and SAPS genes, including *CCL2, IL6, CCL5*, *CCN2* and *TGFB1* (Figure 3l-3r). In parallel, PECs exhibited an activated phenotype, characterized by increased gene expression of *CD44* and *CLDN1*, a tight-junction protein involved in epithelial integrity (Figure 3s and 3t). and increased fibronectin accumulation, detected by immunofluorescence and western blot (Figure 3j and 3k). These data indicate that podocyte-derived senescence signals promote both secondary senescence and PEC activation. As expected, the clearance of senescent podocytes by DQ treatment prevented podocyte-derived secondary senescence in PECs, as evidenced by reduced SA-β-gal activity and normalization of senescence SASP, and PEC activation-related markers (Figure 3). Collectively, these findings identify senescent podocytes as drivers of secondary PEC senescence and reveal a novel senescence-mediated podocyte-to-PEC communication pathway. Importantly, senolytic treatment with DQ effectively interrupts this pathogenic podocyte-to-PEC signaling axis and prevents secondary senescence.

### Therapeutic senolytic treatment improves renal function and attenuates glomerular injury in experimental FSGS

To determine whether senolytic therapy ameliorates established ADR-induced FSGS, DQ was administered after disease onset (Figure 4a). ADR injection resulted in a significant increase in serum creatinine levels at both day 15 and day 22, indicating progressive deterioration of renal function (Figure 4b). Therapeutic administration of DQ significantly reduced serum creatinine levels at both time points, suggesting partial restoration of kidney function. In agreement with previous reports of ADR-induced nephropathy in BALB/c mice ^25^, PAS staining at day 22 revealed severe glomerular injury characterized by podocyte hypertrophy, extracapillary hypercellularity/pseudocrescent formation, and segmental sclerosis (Figure 4c). Histological analysis demonstrated a protective effect of senolytic therapy. Kidneys from ADR+DQ-treated mice exhibited markedly reduced glomerular injury, with preservation of glomerular architecture and absence of segmental sclerosis despite residual extracapillary hypercellularity, resulting in a significant reduction in the glomerular injury score (Figure 4c and 4d). Given the central role of podocyte injury in FSGS, glomerular nephrin expression was assessed by immunofluorescence. ADR mice exhibited a marked reduction in nephrin staining, indicative of podocyte injury, whereas DQ treatment significantly preserved nephrin expression, restoring levels close to those of control animals (Figure 4e and 4f). Consistent with these *in vivo* findings, in cultured human podocytes ADR exposure also reduced nephrin expression, an effect that was significantly prevented by DQ (Figure 4g). Previous studies from our group have implicated the CSF1/CSF1R axis in the pathogenesis of experimental FSGS^30^. In line with these observations, ADR-treated mice kidneys and cultured human podocytes exhibited increased expression of CSF1 and CSF1R, which was markedly downregulated by DQ (Figure 4h and 4i and Supplementary Figure 2a).

**Figure 4.**
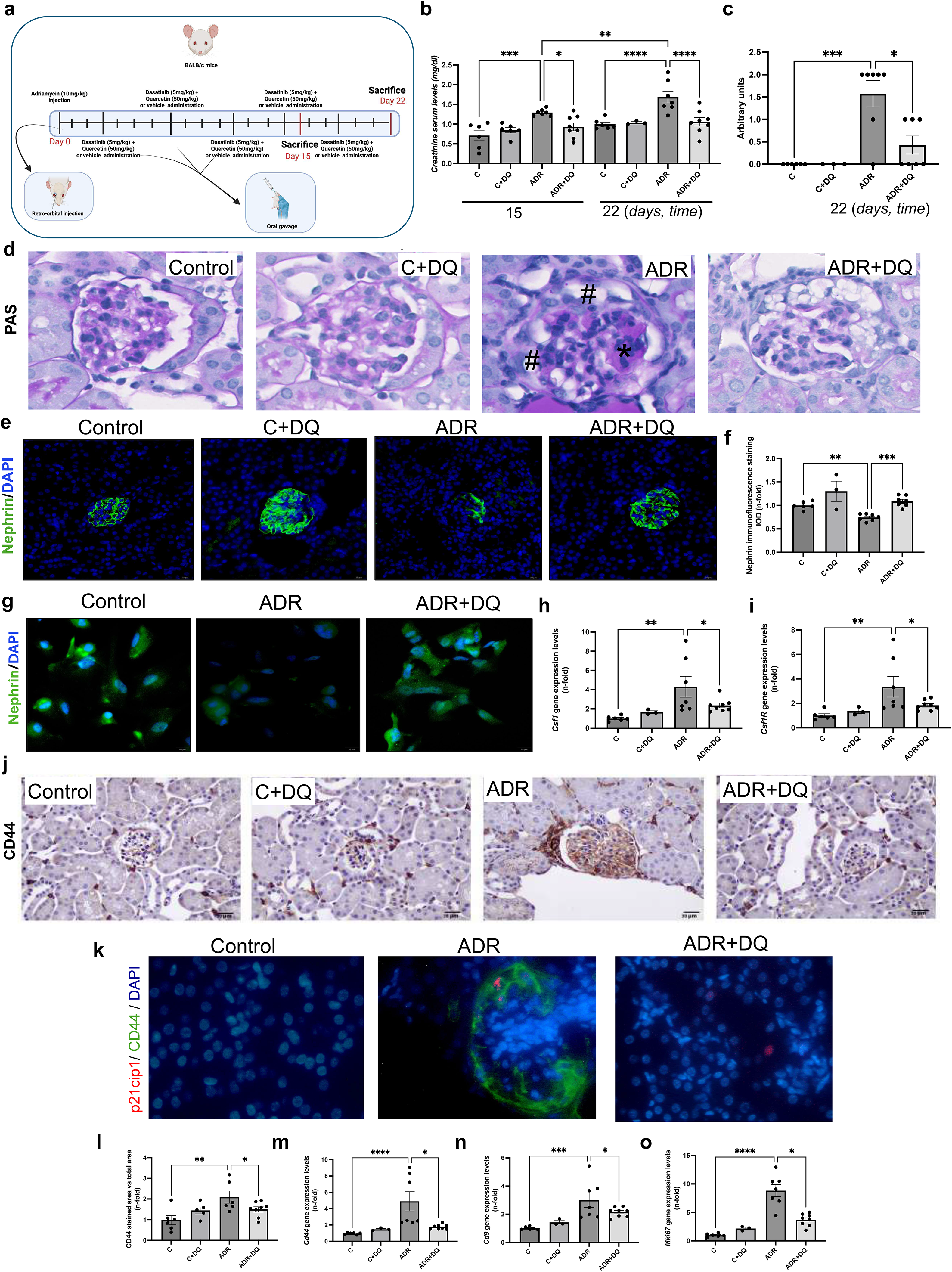
Senolytic therapy ameliorates kidney injury, preserves podocyte integrity and limits parietal epithelial cells (PECs) activation during experimental focal segmental glomerulosclerosis (FSGS). a, Experimental design. Male BALB/c mice received a single retro-orbital injection of adriamycin (ADR; 10 mg/kg) to induce FSGS. The senolytic combination of dasatinib plus quercetin (DǪ; 5 and 50 mg/kg, respectively) was administered by oral gavage starting on day 3 after injury (three doses in the 15-day protocol and five doses in the 22-day protocol). b, Serum creatinine levels in control, ADR-treated and ADR+DǪ-treated mice at days 15 and 22. c, Ǫuantification of glomerular injury scores. d, Representative PAS staining of renal cortex sections at day 22. ADR-treated glomeruli exhibit podocyte hypertrophy, extracapillary hypercellularity/pseudocrescent formation (#) and segmental sclerosis (*), whereas ADR+DǪ-treated mice show reduced glomerular injury and absence of overt segmental sclerosis. e,f, Representative nephrin immunofluorescence (e) and quantification of nephrin-positive area (f) in glomeruli from the indicated experimental groups. (40× magnification). g, Representative nephrin immunofluorescence staining in cultured human podocytes exposed to ADR or ADR+DǪ. h,i, RT–qPCR analysis of *Csf-1* (h) and *Csfr-1* (i) expression in whole-kidney samples. j. Representative CD44 immunohistochemistry in kidney sections from the indicated experimental groups. k Representative immunofluorescence images of p21^Cip1^ (red), the PEC activation marker CD44 (green) and nuclear marker DAPI (blue) in kidneys. l, Ǫuantification of glomerular CD44-positive area. m–o, RT– qPCR analysis of *Cd44*, *CdS* and *MkiC7* expression in whole-kidney samples. Data are presented as mean ± s.e.m. (n=3–8 animals per group). Scale bars, 20 μm unless otherwise indicated. Statistical analyses were performed using ordinary one-way ANOVA followed by Fisher’s LSD test. *p<0.05, **p<0.005, ***p<0.0005.

Because PEC activation is a key driver of FSGS progression, we next examined the therapeutic impact of DQ on this cellular compartment. DQ treatment substantially attenuated glomerular CD44 protein (Figure 4j-4l) and gene expression (Figure 4m) and reduced the number of p21-positive PECs (Figure 4k), indicating a marked suppression of PEC activation and senescence. In line with these histological findings, DQ significantly downregulated the renal expression of the PEC-associated injury markers, *Cd9* and *Mki67,* which were strongly induced following ADR administration (Figure 4n and 4o). No gender differences in response to ADR administration or DQ treatment were found (Supplementary Figure 3). Collectively, these findings demonstrate that therapeutic DQ administration, initiated after disease onset, effectively mitigates renal dysfunction and structural glomerular damage in experimental FSGS, while preserving podocyte integrity and limiting PEC activation.

### Therapeutic DQ treatment reduces glomerular senescence and senescence-associated signaling

ADR-induced FSGS was associated with a marked accumulation of p21^Cip1^-positive cells within the kidney, which was significantly attenuated following DQ administration (Figure 5a and 5b). Additionally, renal expression of senescence-associated cell-cycle regulators, including *Cdkn1a* and *Cdkn2a,* and the DNA damage marker, *H2afX,* was significantly increased in ADR-treated mice and also markedly reduced by senolytic therapy (Figure 5c-5e). These findings suggest that the selected dose and administration schedule of DQ exert renoprotective effects that are mediated, at least in part, by its senolytic activity.

**Figure 5.**
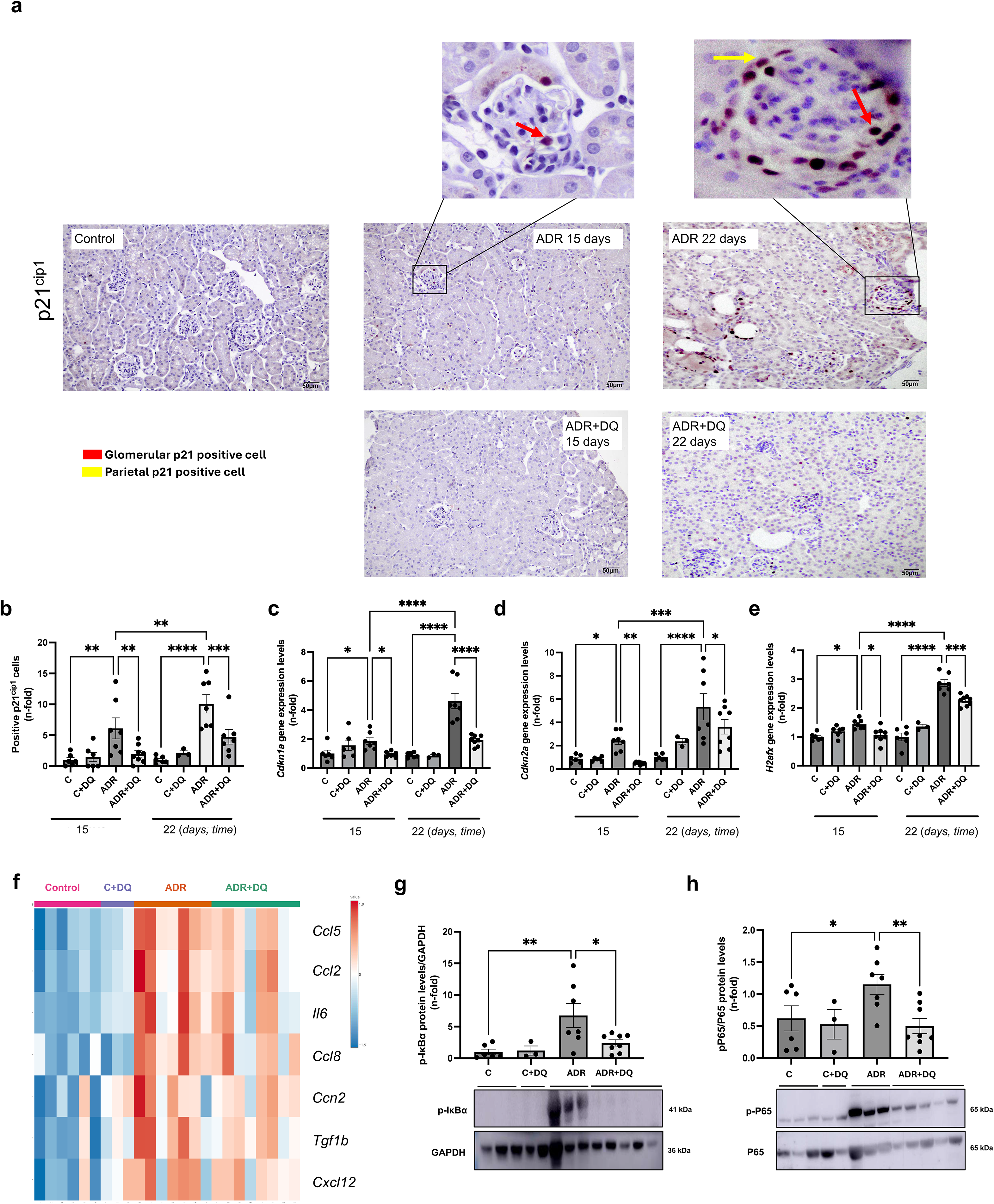
Senolytic therapy reduces glomerular senescence in experimental focal segmental glomerulosclerosis (FSGS). a, Representative p21^Cip1^ immunohistochemistry from control, ADR-treated and ADR+DǪ-treated mice at days 15 and 22. The box shows a magnification of a representative glomeruli. Arrows indicate p21^Cip1^-positive cells in podocytes (red) and PECs (yellow). Scale bars, 50 μm. b, Ǫuantification of p21^Cip1^-positive cells at days 15 and 22, expressed as fold change relative to control mice. p21^Cip1^-positive cell counts were analyzed using the Kruskal–Wallis test followed by Dunn’s multiple-comparison test. c–e, RT–qPCR analysis of the senescence-associated cell-cycle arrest genes *Cdkn1a*, *Cdkn2a* and *H2afX* in kidneys from the indicated experimental groups. f, Heat map showing relative expression of senescence-associated secretory phenotype (SASP) genes in kidneys from different groups. Heatmap represents the row *Z*-score of the relative expression levels of inflammatory markers. Relative gene expression was analyzed by RT-qPCR from total RNA of renal tissue, using *Gapdh* as housekeeping gene. **Activation of NF-κB in FSGS is prevented by DǪ treatment**. Representative immunoblots and densitometric quantification of phosphorylated IκBα (g) and p65 NF-κB (h) in kidneys from control, ADR-treated and ADR+DǪ-treated mice. Data are presented as mean ± s.e.m. (n=3–8 animals per group). Each symbol represents one mouse. Statistical analyses were performed using ordinary one-way ANOVA followed by Fisher’s LSD test. *p<0.05, **p<0.005, ***p<0.0005.

One of the main features of senescent cells is the acquisition of a characteristic secretome, the SAPS, which is responsible for the spread of secondary senescence. Therefore, we also evaluated whether experimental FSGS was associated with upregulation of key SASP components. ADR mice exhibited increased renal expression of *Ccl5, Ccl2, Il6, Ccl8, Ccn2, Tgfb1* and *Cxcl12*, which was markedly attenuated by DQ treatment (Figure 5f). Likewise, senescent cultured human podocytes showed increased expression of SASP components, including the pro-inflammatory and pro-fibrotic mediators *CCL2, CCL5, IL-6, CXCL12, CCN2* and *TGFB1,* whereas DQ effectively reduced their expression (Supplementary Figure 2b).

Next, we examined signaling pathways involved in the senescence program. Since the canonical NF-κB pathway is a major regulator of SASP gene expression^16^, we assessed NF-κB activation *in vivo* and *in vitro*. ADR-treated kidneys displayed increased phosphorylation of IκBα and p65 NF-κB, demonstrating NF-κB pathway activation. Notably, therapeutic administration of DQ markedly attenuated these changes (Figure 5g and 5h). Furthermore, NF-κB activation was also observed in ADR-induced podocyte senescence and was also effectively prevented by DQ treatment (Supplementary Figure 2c). ERK and STAT3 signaling have also been implicated in the regulation of senescence-associated transcriptional programs and SASP^31^. ADR exposure increased ERK and STAT3 phosphorylation, whereas DQ markedly attenuated activation of both kinases in cultured human podocytes (Supplementary Figure 2d and 2e).

Defective autophagy is another recognized feature of senescent cells^32^. Thus, we evaluated some autophagy-associated pathways. In ADR-induced FSGS, we observed reduced LC3-II levels and accumulation of p62, which were reverted by DQ (Supplementary Figure 4a and 4b). Similarly, renal expression of the autophagy-related genes *Atg5, Becn1*, and *Atg7* was dysregulated in ADR-treated kidneys and normalized following senolytic therapy (Supplementary Figure 4c-4e), suggesting a restoration of autophagy-associated homeostasis following DQ treatment.

Senescent cells evade apoptosis through activation of senescent cell anti-apoptotic pathways (SCAPs), including members of the BCL-2 family^16,31^. ADR-induced injury was associated with increased renal *Bcl2l* expression, which was significantly reduced by DQ treatment (Supplementary Figure 4f). Given the established role of SRC kinases in cell survival^33^ and their recognition as major targets of dasatinib^34,35^, we next examined SRC activation in human podocytes. ADR exposure increased SRC phosphorylation, whereas DQ markedly attenuated SRC activation (Supplementary Figure 4g). Together, these findings support the ability of DQ to disrupt prosurvival programs associated with cellular senescence.

Taken together, these results demonstrate that therapeutic DQ administration mitigates the senescence program activated in experimental FSGS. Beyond reducing the accumulation of senescent cells, DQ suppressed SASP production, inhibited key senescence-associated signaling pathways, restored autophagy-related mechanisms, and impaired SCAP-dependent survival signaling, supporting a central role for cellular senescence in disease progression and therapeutic response.

### Proteomics Identifies Changes in Proteins Linked to Podocyte Injury in FSGS that are restored by DQ

To further investigate novel mediators and molecular pathways underlying the beneficial effects of senolytic therapy in FSGS, untargeted proteomic profiling was performed on kidney samples obtained from mice with early-stage ADR-induced FSGS (day 15). On average, 5,640 proteins were identified with high confidence (FDR ≤1%), of which 5,626 were quantified across groups. The comparison of the ADR group with controls identified 424 proteins differentially expressed at nominal significance (p<0.05), of which 25 remained significant after FDR correction (FDR<0.05), whereas the comparison of ADR+DQ with ADR groups identified 270 differentially expressed proteins (p<0.05), of which only five remained significant after FDR correction (Figure 6a). Proteomic profiling revealed significant alterations in proteins associated with podocyte injury. Among these differentially expressed proteins G protein-coupled receptor class C group 5 member A (GPRC5A), a podocyte-specific protein^36^, and SH3-domain binding protein 5 (SH3BP5) were markedly downregulated in injured kidneys and restored by DQ treatment (Figure 6a). The translational relevance of these findings was confirmed by searching publicly available human FSGS transcriptomic datasets using Nephroseq v5. As observed in preclinical FSGS, glomerular expression of *GPRC5A* was downregulated in FSGS and more specifically in collapsing FSGS (from data set^37^) (Figure 6b and 6c). Regarding SH3BP5 gene diminution of its levels were found in collapsing FSGS (from data set ^37^) and nephrotic syndrome patients (from data set ^38^) (Figure 6d and 6e).

**Figure 6.**
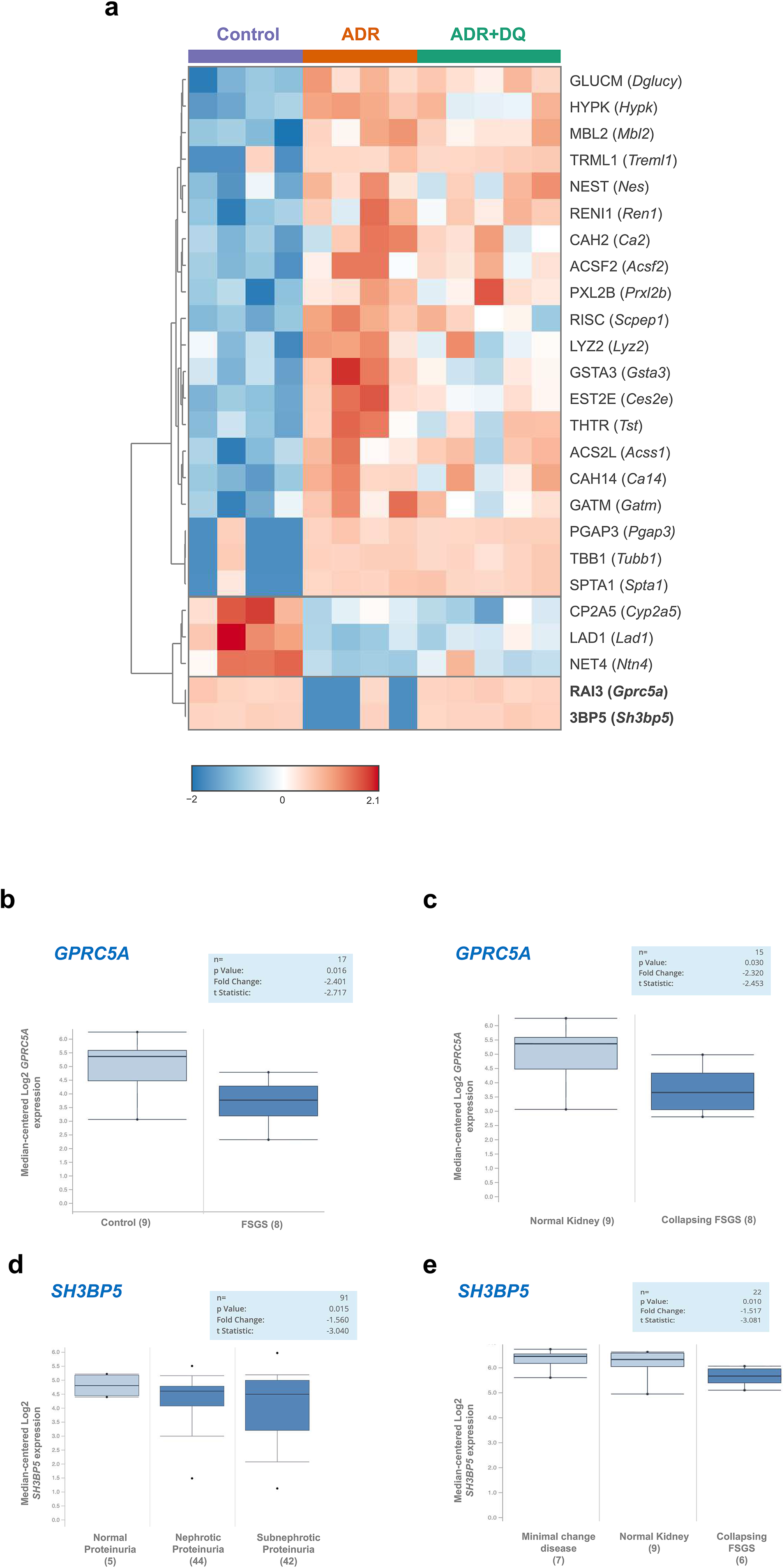
Proteomic profiling reveals restoration of proteins depleted in experimental FSGS by senolytic therapy. a, Heatmap showing proteins (and their corresponding genes) differentially expressed among control, ADR-treated and ADR+DǪ-treated mice at day 15. Displayed proteins were differentially abundant following false discovery rate correction (FDR < 0.05). Color intensity indicates relative protein expression levels. GPRC5A and SH3BP5 were among the proteins reduced in ADR-treated mice and restored by senolytic treatment. **Human FSGS datasets validate the reduced expression of GPRC5A and SH3BP5 identified in experimental FSGS.** Human transcriptomic datasets were interrogated using Nephroseq v5. b,c, Glomerular *GPRC5A* gene expression in the Hodgin FSGS dataset^37^. b, Primary FSGS versus healthy controls. c, Collapsing FSGS versus healthy controls. d,e, Glomerular *SH3BP5* expression in the Mariani Nephrotic Syndrome Glom (NEPTUNE) dataset^38^ (GSE197307) and the Hodgin FSGS dataset^37^. Both genes were significantly downregulated in patients with FSGS compared with healthy controls, consistent with the proteomic alterations observed in experimental FSGS. Box plots show median-centered log2 expression values; P values, fold changes and t-statistics are indicated within each panel.

### Proteomic Profiling Identifies Interferon and STAT3 Signaling as Key Targets of Senolytic Therapy in FSGS

Functional analysis by Gene Set Enrichment Analysis (GSEA) revealed significant enrichment of the interferon-γ response, interferon-α response, IL6-JAK-STAT3 signalling, and allograft rejection pathways in ADR-treated kidneys (Figure 7a and 7b). Interferon-α and interferon-γ response signatures were no longer enriched in ADR+DQ kidneys compared with controls, suggesting attenuation of interferon-related signalling following senolytic treatment. Moreover, protein-protein interaction analysis of the core enrichment proteins identified a densely interconnected inflammatory network enriched in interferon-responsive and STAT3-associated proteins (Figure 7c). These proteomic findings were validated at gene levels by qPCR analysis. ADR mice presented increased renal expression of *Ifna, Ifng, Ifit1, Mx1, Isg15, Oasl2,* and *B2m*, which was significantly reduced following DQ administration (Figure 7d). Moreover, ADR upregulated interferon-genes in cultured human podocytes, including *IFNI44, IFIT1, IFI27,* and *ISG15*, whose expression was significantly inhibited by DQ (Supplementary Figure 5). These data indicate that DQ suppresses a coordinated interferon-driven inflammatory program associated with disease progression. Moreover, the evaluation of publicly available human FSGS databases nephroseq V5 showed that in the European Renal cDNA Bank (ERCB) cohort^39^ (Ju CKD Glom) the expression of multiple interferon-stimulated genes (ISGs), including *ISG15, IRF7, IRF1, CXCL10, IFI44* and *IFIT1* (Supplementary Figure 6), was significantly increased in patients with FSGS compared with healthy controls.

**Figure 7.**
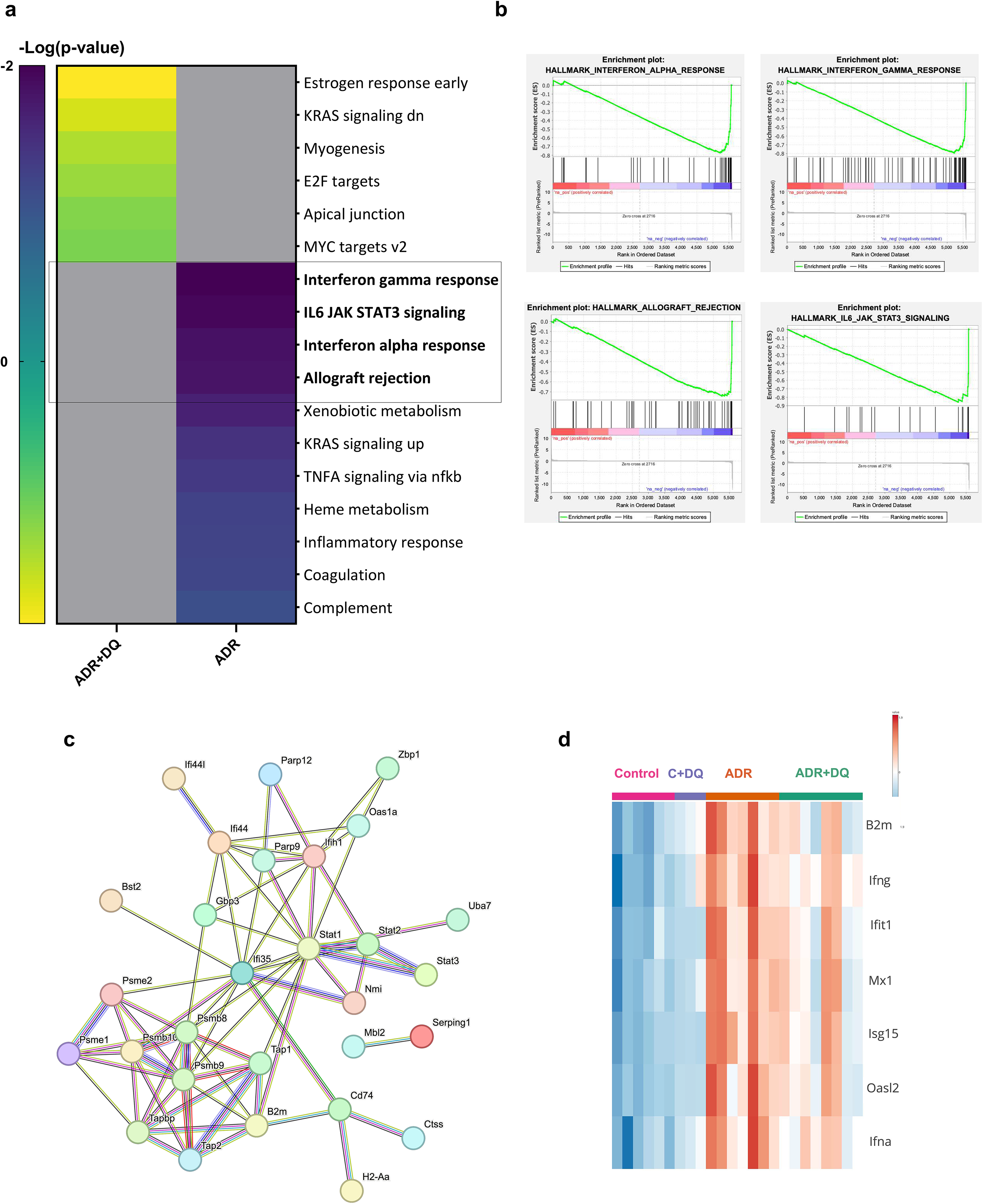
Proteomic profiling reveals suppression of interferon- and signaling by senolytic therapy in experimental FSGS. a-b, Gene Set Enrichment Analysis (GSEA) of proteomic data comparing ADR+DǪ-treated mice with ADR-treated mice at day 15. a, Heatmap showing enriched gene sets, highlighting four pathways negatively enriched in ADR+DǪ-treated mice: interferon-γ response, IL6-JAK-STAT3 signaling, interferon-α response and allograft rejection. b, GSEA enrichment plots for the four highlighted pathways. Negative enrichment curves shifted to the right (green line), together with the distribution of core enrichment proteins within the ranked dataset (black vertical lines), indicate preferential enrichment in ADR-treated compared with ADR+DǪ-treated mice. c, STRING protein–protein interaction network of core enrichment proteins from the four pathways modulated by DǪ treatment. The network was generated using evidence-based interactions with a high-confidence interaction score of 0.700. d, Heatmap showing qPCR validation of the selected genes (*Ifna, Ifng, Ifit1, Mx1, Isg15* and *Oasl2* and *B2m*) in kidneys from control, control+DǪ, ADR-treated and ADR+DǪ-treated mice. Gene expression data are presented as mean ± s.e.m. (n = 3–8 animals per group). Statistical analyses were performed using ordinary one-way ANOVA followed by Fisher’s LSD test. *p<0.05, **p<0.005, ***p<0.0005

## Discussion

Our findings identify senescence propagation within the glomerulus as a previously unrecognized mechanism of FSGS progression. Senescent podocytes promote secondary senescence in neighboring PECs through inflammatory signaling networks involving NF-κB, interferon, and STAT3, while senolytic therapy effectively counteracts this pathogenic cascade. These observations support a model in which cellular senescence actively orchestrates the progression of glomerular injury through pathogenic intercellular communication.

Podocyte depletion is widely recognized as the initiating event in FSGS^1,3,16^; however, the mechanisms linking podocyte injury to progressive glomerulosclerosis remain incompletely understood. A major finding of the present study is the identification of senescent PECs in both human and experimental FSGS. While podocyte senescence has previously been implicated in glomerular injury^17,28,40^, the presence of senescent PECs suggests that senescence extends beyond a single glomerular cell type and involves the broader glomerular niche. In our preclinical FSGS model, temporal analyses demonstrated that senescent podocytes appeared early during disease progression, whereas senescent PECs emerged only at later stages. Importantly, *in vitro* conditioned-medium experiments showed that senescent podocytes induce a senescent phenotype in PECs, providing direct evidence for secondary senescence within the glomerular compartment. Recent single-cell and single-nucleus transcriptomic studies demonstrate extensive crosstalk between podocytes and PECs^41,42^, supporting a central role for PECs in glomerular disease progression. Podocyte loss remodels the signaling milieu of Bowman’s space, including NOTCH and WNT/β-catenin activation^43,44^, activation of NF-κB pathway ^45^, CXCL12-dependent podocyte-progenitor feedback^46^ and CSF1/CSF1R axis ^30^. Our findings extend this intercellular signaling network by identifying cellular senescence as a mechanism through which senescent podocytes can promote a senescence-associated response in neighboring PECs via SASP-related inflammatory signaling involving NF-κB and interferon/STAT3 pathways. IL-6 may be particularly relevant in this context given its ability to activate STAT3 signaling^47^, whereas the CCL2-CCR2 axis has been implicated in inflammatory cell recruitment and disease progression in experimental FSGS^48^. Together, our preclinical findings support a model in which senescent podocytes reshape the glomerular microenvironment through SASP-associated mediators, promoting senescence in neighboring PECs and thereby amplifying glomerular injury.

Pharmacological targeting of cellular senescence with the senolytic combination DQ markedly attenuated established experimental FSGS, improving kidney function and reducing glomerular injury. DQ preserved the podocyte phenotype, as reflected by restored nephrin expression in ADR-injured kidneys and by maintenance of GPRC5A, a podocyte-enriched protein whose loss increases susceptibility to glomerular injury^36^. Importantly, DQ reduced the senescent cell burden and SASP-associated inflammatory responses and prevented the propagation of secondary senescence to PECs. Our *in vitro* studies demonstrated that DQ suppressed NF-κB, ERK, and STAT3 activation in ADR-induced senescent podocytes and experimental FSGS, together with downregulation of SASP-related genes, including IL-6 and CCL2. These findings are consistent with studies in experimental diabetic kidney disease showing that DQ improves kidney function and reduces cellular senescence and inflammation, while preserving podocyte differentiation^24,49^, suggesting that senescence-associated mechanisms may represent shared drivers of progressive kidney injury.

PECs may represent a critical target of this senescence-associated signaling, as their activation contributes to sclerotic lesion formation and FSGS progression^50^. DQ markedly reduced PEC activation and secondary senescence, as indicated by decreased levels of CD9, CD44, and proliferative markers and fewer p21-positive PECs, as well as by suppression of the CSF1/CSF1R axis, previously identified by our group as a key driver of PEC activation in FSGS^30^. In addition, experimental FSGS was associated with increased expression of the profibrotic mediators CCN2 and TGF-β1, which may couple tissue remodeling to the induction and maintenance of cellular senescence and ECM accumulation^51,52^. Together, our findings suggest that senescent podocytes may act as signaling hubs that amplify glomerular injury through SASP-driven inflammation and secondary PEC senescence, while senolytic therapy can interrupt this maladaptive podocyte–PEC crosstalk to limit progression to segmental glomerulosclerosis.

The beneficial effects of DQ likely involve disruption of SCAPs, which allow senescent cells to persist despite chronic cellular stress and DNA damage^53^. In this context, SRC family kinases contribute to both cell survival and inflammatory signaling^33^ and represent major molecular targets of dasatinib^34,35^. In line with this mechanism, DQ reduced SRC phosphorylation and restored *Bcl2l* gene deregulation, suggesting destabilization of survival programs that sustain senescent podocytes and perpetuate glomerular injury. Multi-tissue analyses in aged mice have shown that DQ remodels immune responses, reduces tissue inflammation, and attenuates multiple hallmarks of aging in a tissue-specific manner ^54^, suggesting that the benefits of senolytic interventions extend beyond senescent cell clearance, as reported here. The therapeutic potential of senolytic interventions is increasingly supported by early clinical studies. Intermittent treatment with DQ improved physical performance in idiopathic pulmonary fibrosis^55,56^ and reduced senescent cell burden and circulating SASP factors in diabetic kidney disease^56^, providing proof-of-concept that senescence can be therapeutically targeted in humans. The growing number of clinical trials across neurodegenerative, metabolic, musculoskeletal, and renal disorders^57^ further highlights the broad translational potential of senolytic therapies and supports their investigation in FSGS. Proteomic profiling identified interferon signaling as one of the most prominent pathways dysregulated in experimental FSGS. This finding was subsequently validated through complementary gene expression analyses in injured podocytes, experimental FSGS, and independent human transcriptomic datasets, where several interferon-pathway components, including ISG15 and IFIT1, were consistently upregulated. The conservation of this molecular signature across experimental models and human disease supports a pathogenic role for interferon-dependent pathways in senescence-associated inflammation and progressive glomerular injury. Importantly, DQ normalized interferon-responsive gene expression *in vitro* and *in vivo*, suggesting a novel mechanism underlying its beneficial effects. Interestingly, a recent study identifying universal transcriptomic hallmarks of mammalian ageing and mortality identified interferon signaling among the pathways negatively associated with lifespan^26^. Our findings extend the relevance of interferon-associated signatures to a disease setting, while underscoring the importance of their biological context. In FSGS, we demonstrate that senescent cells contribute to disease progression and that this phenotype is accompanied by activation of interferon signaling, which is attenuated by senolytic treatment. These findings suggest that interferon signaling contributes to the senescence-associated response in FSGS and may promote glomerular injury. They also underscore the context-dependent nature of cellular senescence, as individual markers and pathways may have different relevance across cell types and between physiological ageing and chronic disease. Understanding these differences will be important for defining disease-specific senescence signatures and, ultimately, for developing senescence-targeted therapies. The activation of interferon signaling in FSGS also has important translational implications, as clinically available therapies targeting the interferon pathway provide an opportunity to therapeutically modulate this senescence-associated response^58^. These include blockade of type I interferon signaling with the anti-IFNAR1 monoclonal antibody anifrolumab ^59^, as well as inhibition of downstream JAK-STAT signaling using JAK inhibitors^60^. Collectively, these findings position interferon signaling as a potential mechanistic driver of FSGS progression and support the evaluation of interferon- and JAK/STAT-targeted interventions as therapeutic strategies to slow disease progression.

In summary, our findings suggest that senescence spreads from injured podocytes to PECs, amplifying glomerular damage and sclerosis. Importantly, therapeutic senolysis with DQ reduced senescent cell burden, improved renal function, and attenuated proteinuria and glomerular injury in experimental FSGS. By identifying cellular senescence as a key driver of disease progression, our results provide a compelling rationale for exploring senolytic, as well as interferon, therapies as novel potential treatment strategy for patients with progressive FSGS.

## Data Statement

The mass spectrometry proteomics data have been deposited to the ProteomeXchange Consortium via the PRIDE [PubMed ID: 39494541] partner repository with the dataset identifier PXD081301. Other data available in the main text or in the Supplementary Material will be made accessible by the authors on request.

## Submission details

**Project Name:** Glomerular Senescence in Focal Segmental Glomerulosclerosis Represents a Therapeutic Target of Dasatinib Plus Quercetin

**Project accession:** PXD081301

**Project DOI:** Not applicable

## Acknowledgements

The results of this work have been previously presented at several scientific meetings, including the 63rd ERA Congress, Glasgow, UK, Annual Meeting of the Madrid Society of Nephrology (SOMANE) 2026 in La Granja, Spain. Meeting of the Spanish Society of Nephrology SENEFRO2026 in Murcia, Spain and European Renal Cell Study Group Meeting 2026 Paris, France.

## Author contributions

A. B-V, S.R-M and M.R-O conceived and designed the work. A. B-V, C. G-C, T.Q.N, F.M.S., S.C, N.M-B, A.S and S.R-M performed the experiments and obtained the data. A. B-V., M.R-O, C. G-C, T.Q.N, F.M.S analyzed and interpreted the results. A. B-V; S.R-M and M.R-O drafted the manuscript. A.S, C. G-C, S.R-M, A.O F.M.S., and M.R-O edited and revised the manuscript. All authors approved the final version of the manuscript.

---

**Supplementary Figure 1.**
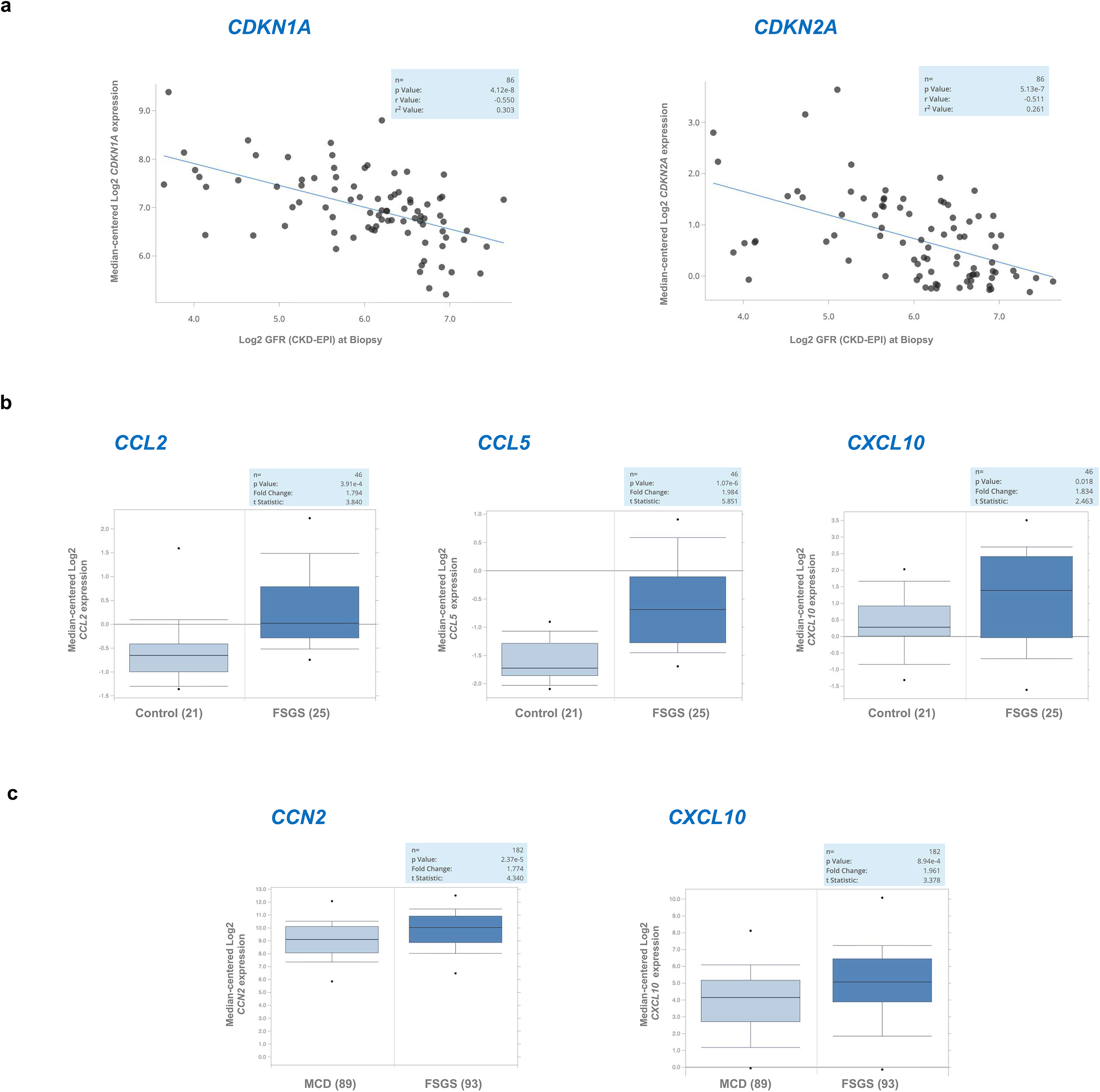
Evidence of senescence-associated pathways in human focal segmental glomerulosclerosis (FSGS) transcriptomic datasets. a. Glomerular expression of the senescence markers p21^Cip1^ (*CDKN1A)* and p16^INK4a^ (*CDKN2A*) was analyzed in the Mariani Nephrotic Syndrome Glom RNA-seq dataset^38^ (GSE197307) using the Nephroseq v5. Scatter plots show the relationship between gene expression and estimated glomerular filtration rate (eGFR) in patients with FSGS (n = 86), revealing an association between higher expression of senescence markers and lower eGFR. Lines represent linear regression fits. b, Analysis of independent human FSGS transcriptomic datasets revealed increased glomerular expression of senescence-associated secretory phenotype (SASP) mediators. In the Ju CKD Glom dataset ^39^ (GSE104948), comprising microdissected glomeruli from the European Renal cDNA Bank (ERCB) cohort, *CCL2, CCL5,* and *CXCL10* expression was significantly increased in FSGS compared with healthy living donors. c. Similarly, analysis of the Mariani Nephrotic Syndrome Glom RNA-seq dataset showed higher glomerular expression of *CXCL10* and *CCN2* in FSGS than in minimal change disease (MCD). Box plots show median-centered log2 expression values. Fold changes, t-statistics, and P values are indicated in each panel.

**Supplementary Figure 2.**
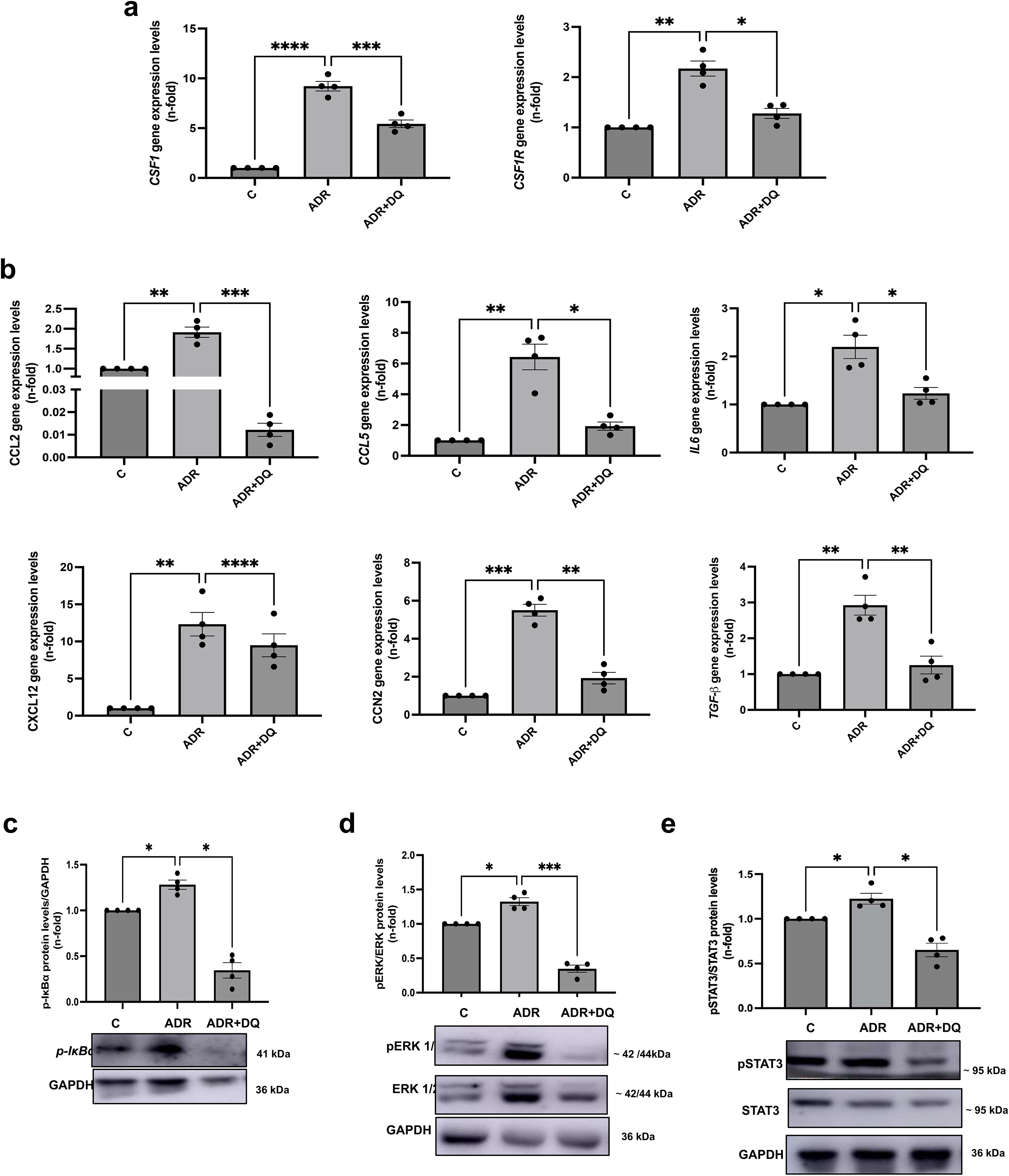
Dasatinib plus quercetin (DǪ) treatment suppresses Adriamycin-induced gene upregulation in cultured human podocytes. Podocytes were exposed to ADR (1 μg/mL, 1 or 24 h) with or without treatment with Dasatinib 600nM and Ǫuercetin 30uM (DǪ). Gene expression was evaluated by RT–qPCR analysis. a, Modulation of the CSF1/CSF1R pathway by DǪ treatment. b, Expression of senescence-associated secretory phenotype (SASP) mediators, including CCL2, CCL5, IL6, CXCL12, CCN2 and TGFB1. Representative immunoblots and densitometric quantification of phosphorylated IκBα (i) ERK (j) and STAT-3 (k). Data are presented as mean ± s.e.m. from 3–4 independent experiments. Statistical analyses were performed using one-way ANOVA followed by Holm–Šídák multiple-comparison testing. *p<0.05, **p<0.005, ***p<0.0005.

**Supplementary Figure 3.**
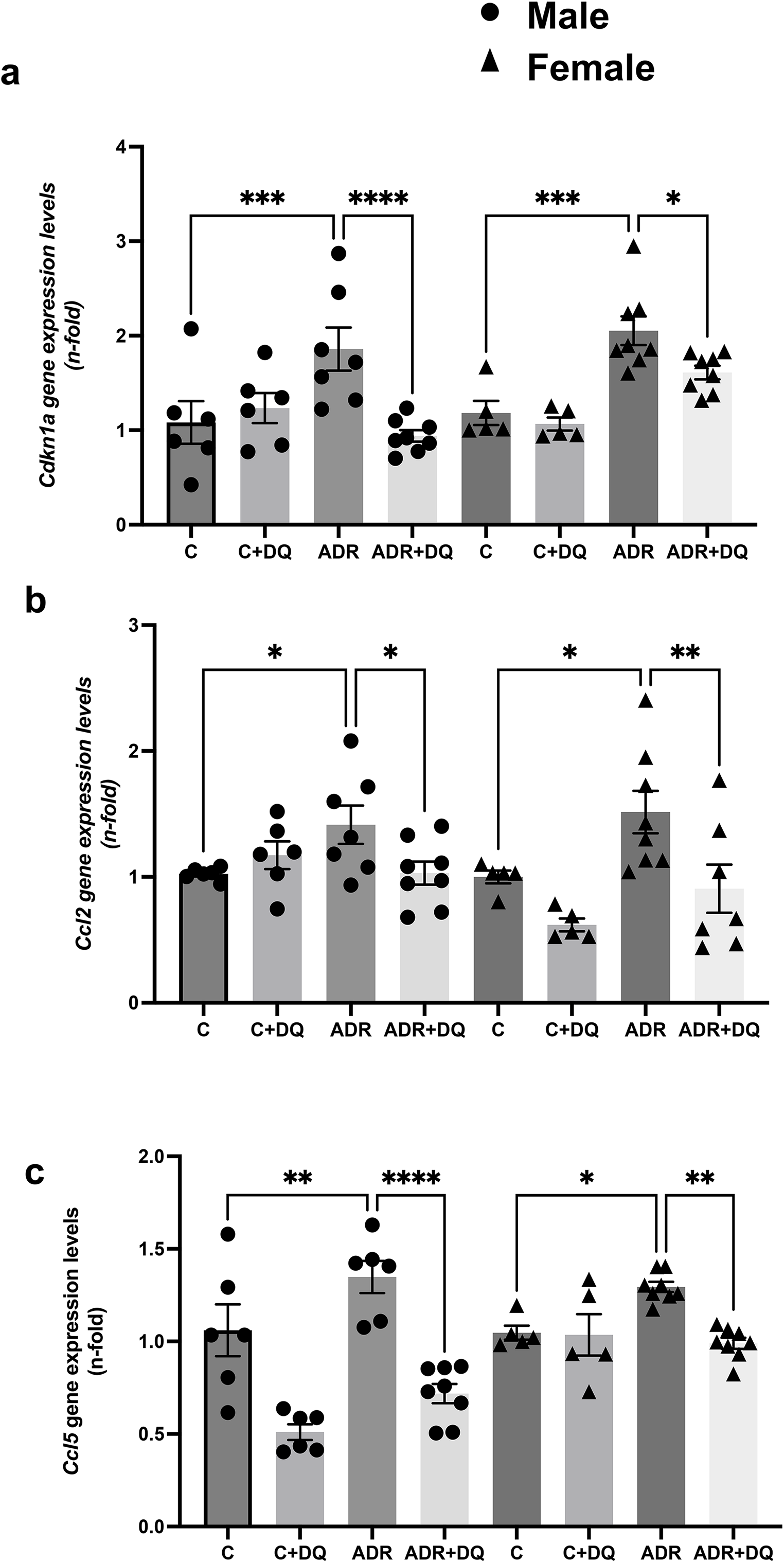
Female mice recapitulate the response to senolytic DǪ therapy in ADR-induced FSGS. Female BALB/c mice received a single retro-orbital injection of adriamycin (ADR; 10 mg/kg) to induce FSGS. Senolytic combination dasatinib plus quercetin (DǪ 5 and 50 mg/kg, respectively) was administered via oral gavage starting on day 3 post-injury (three doses) and mice were analysed on day 15. RT–qPCR analysis of *Cdkn1, ccl2* and *ccl5* expression in kidneys from the indicated experimental groups. Each symbol represents one mouse. (n=5–8 animals per group). Statistical analyses were performed using ordinary one-way ANOVA followed by Fisher’s LSD test. *p<0.05, **p<0.005, ***p<0.0005.

**Supplementary Figure 4.**
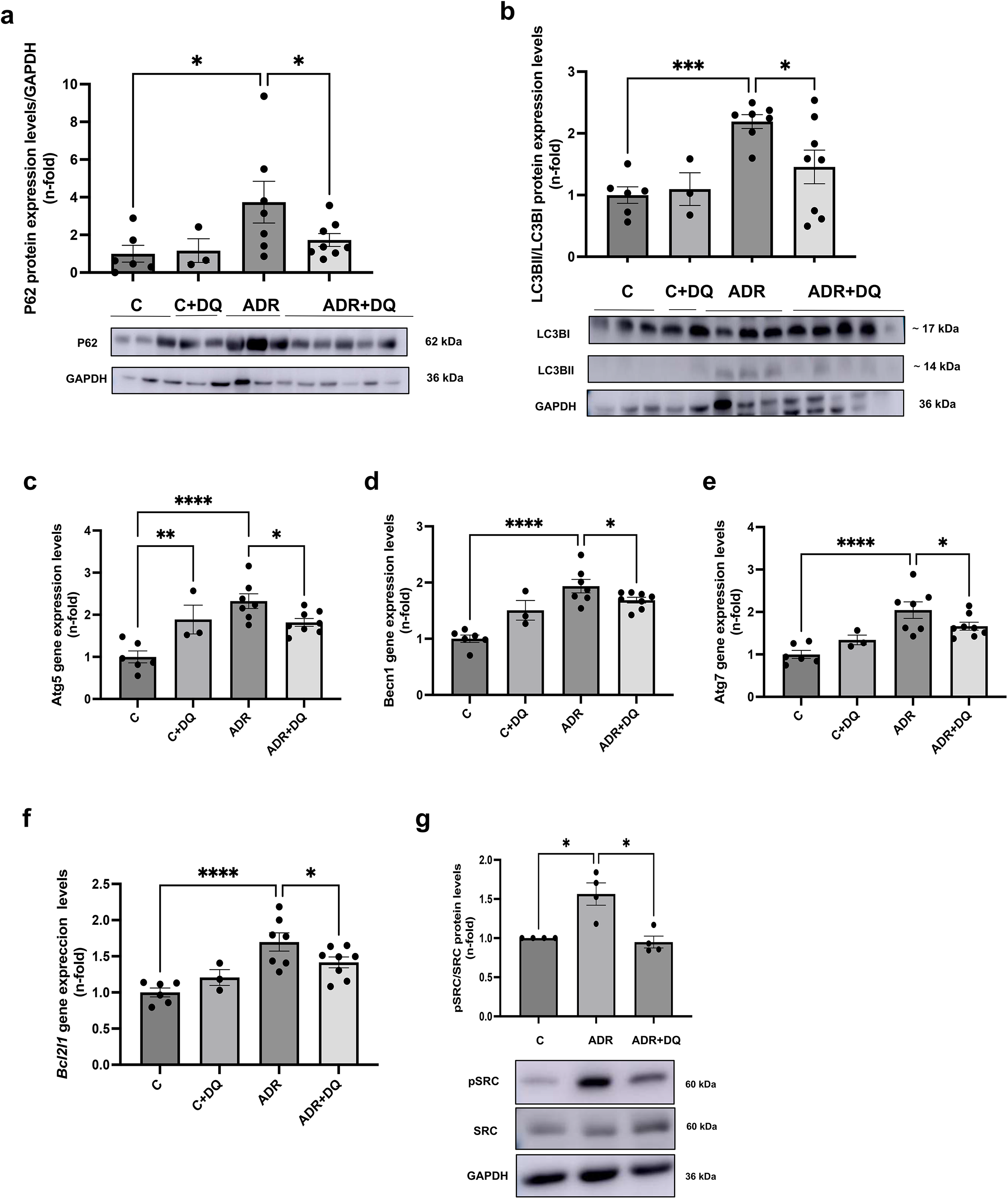
Senolytic therapy restores autophagy and apoptosis-related pathways in experimental FSGS. a,b, Representative immunoblots of LC3-II (a) and p62 (b) expression in kidneys from control, ADR-treated and ADR+DǪ-treated mice and its corresponding densitometric quantification. c–f, RT–qPCR analysis of *Atg5* (c), *Becn1 (d) Atg7* (e) and *Bcl2l* (f) expression in kidneys from the indicated experimental groups. g. Representative immunoblots of phosphorylated SRC (p-SRC) expression in kidneys and its corresponding densitometric quantification. Data are presented as mean ± s.e.m. (n = 3–7animals per group). Each symbol represents one mouse. Statistical analyses were performed using one-way ANOVA followed by Fisher’s LSD test. *p<0.05, **p<0.005, ***p<0.0005.

**Supplementary Figure 5.**
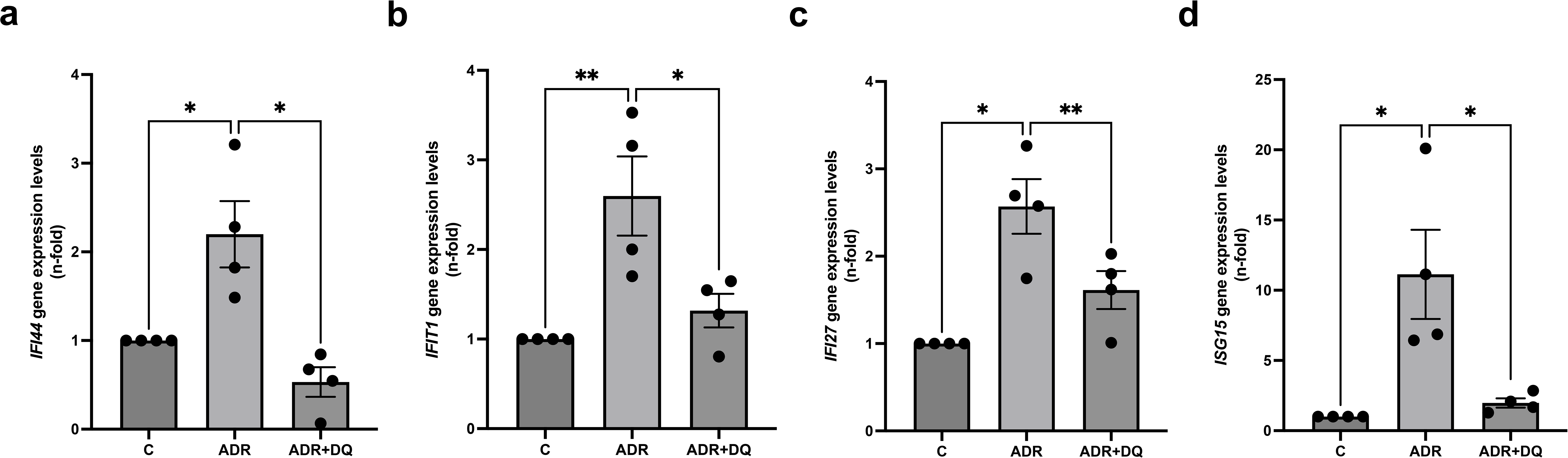
Interferon-related genes in cultured human podocytes. Human podocytes were exposed to ADR (1 μg mL⁻¹, 24 h) in the presence or absence of dasatinib (600 nM) and quercetin (30 μM) (DǪ). RT-qPCR analysis of interferon-genes including *IFI44, IFIT1, IFI27* and *ISG15.* Data are presented as mean ± s.e.m. from 3–4 independent experiments. Statistical analyses were performed using one-way ANOVA followed by Holm–Šídák multiple-comparison testing. *p<0.05, **p<0.005, ***p<0.0005.

**Supplementary Figure 6.**
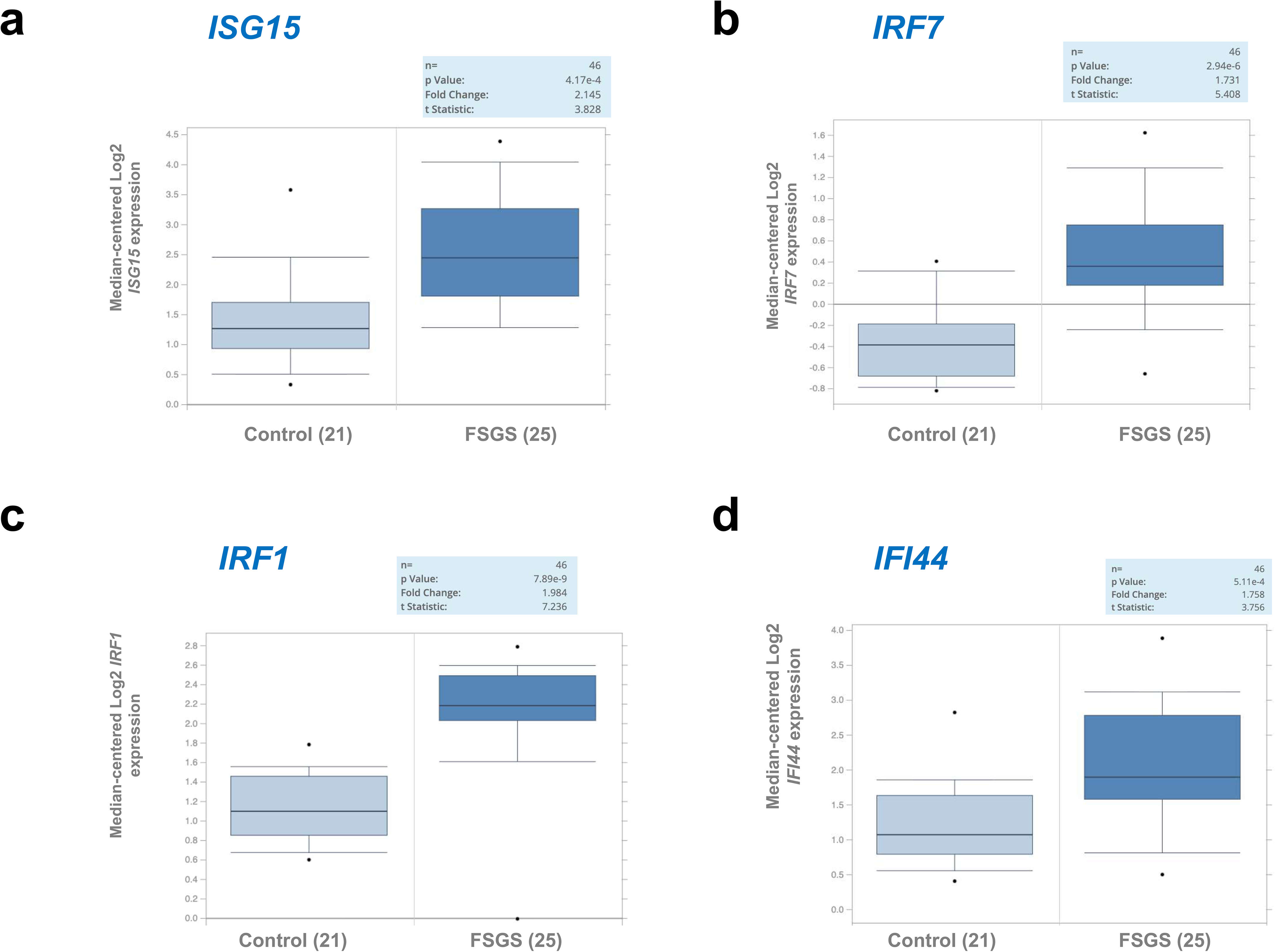
Human FSGS transcriptomic datasets reveal upregulation of interferon-responsive genes. Glomerular expression of *ISG15, IRF7, CXCL10* and *IFIT3* was analyzed using Nephroseq v5 in the European Renal cDNA Bank (ERCB) cohort (Ju CKD Glom^39^; GSE104948), comprising microdissected glomerular samples from patients with FSGS and healthy living donors. Gene expression was assessed using the Affymetrix GeneChip Human Genome U133 Plus 2.0 platform. Box plots show median-centered log2 expression values. Fold changes, t-statistics and P values for the comparison between FSGS and healthy living donors are indicated within each panel.

