## Supplemental Methods for "Podocyte-to-Parietal Epithelial Cell Senescence Drives Experimental Focal Segmental Glomerulosclerosis and is reversed by Dasatinib Plus Quercetin"

**Supplementary Material and Methods**

**Human samples**

Six human kidney biopsies with FSGS were selected from the pathology archives of UMC Utrecht. All experiments on human samples followed all relevant guidelines and regulations. The use of kidney biopsies for research purposes was approved by the Biobank Research Ethics Committee of UMC Utrecht, Netherlands (TCBio 19-945 269). The studies were conducted in accordance with the local legislation and institutional requirements. All patients provided written informed consent and the study was performed in accordance with the Declaration of Helsinki.

**Cell cultures**

*Human podocytes*

The cell line is conditionally immortalized through a temperature-sensitive SV40 large T-antigen transgene, as previously reported [1]. After shifting the culture temperature from 33 °C to 37 °C, the cells complete differentiation within 14 days. They are maintained in RPMI-1640 medium (Lonza, Leeds, UK), supplemented with 10% fetal bovine serum (FBS, A5256701, Gibco), 0.1% L-glutamine (G7513, Sigma-Aldrich), 0,1% insulin–transferrin–selenium (ITS, 41400-045, Gibco, Sigma-Aldrich,) and 0,1% Zhelsield (13-0050, MinervaBiooLabs, Germany) until they have been used. The day before the experiment cells were supplemented with 2% fetal bovine serum without extracellular vesicle for 24h, then cells were exposed to DMSO 0,1%, Adriamycin (ADR, D1515, Sigma-Aldrich) 1µg alone or in the presence of the senolytic combination Dasatinib and Quercetin (DQ) (at doses Dasatinib (D); 600nM; SML2589, Sigma-Aldrich; Quercetin (Q); 30µM Q4951, Sigma-Aldrich).

*Human parietal epitelial cells*

Human PECs (hPECs) were kindly provided by Dr. Catherine Meyer-Schwesinger (UKE, Hamburg)[2]. The cells were maintained in a 1:1 mixture of RPMI 1640 medium with L-Glutamine and 25 mM HEPES (22400-089, Gibco) and endothelial cell basal medium (EBM, CC-312 Lonza), supplemented with 15% FBS, 0.1% gentamicin/amphotericin B (L0012, Cultek/ 30-003-CF, Corning), 2 mM L-glutamine, 100 U/ml penicillin/streptomycin, 5 µg/ml ITS, 0.1% hydrocortisone (H0888, Sigma-Aldrich) and 100 mM Sodium Pyruvate (11360-070, Gibco), 0.4% bovine brain extract (BBE, CC-4092, Lonza), and 0.1% human epidermal growth factor (hEGF, CC-4107 Lonza). Cultures were maintained at 33 °C in 5% $CO_2$, and differentiation was induced by shifting the temperature to 37 °C for 15 days prior to experimental use. Once ready, cells were exposed to podocyte-derived conditional media (PDCM).

*Primary culture renal cells*

Primary murine podocyte and PEC (PECs) were isolated from the kidney of BALB/c mice as previously described[3]. Renal cortex fragments were digested with 0.5 mg/ml collagenase type II

(Sigma, C6885) at 37 °C for 15 minutes. Following enzymatic digestion, the tissue was mechanically homogenized by passing it through a 100 µm cell strainer. Red blood cells were lysed for 10 min at 4ºC. The cells suspension was centrifugated at 1500 rpm for 10 min and the supernatant was discarded. The resulting pellet was resuspended in phosphate-buffered saline (PBS) and passed through a 40 µm cell strainer to separate tubular cells from encapsulated glomeruli. For murine PEC cultures, isolated glomeruli were maintained in RPMI 1640 medium supplemented with 10% FBS, 0,1% L-glutamine, 0,1% ITS and 0,1% 100 U/ml penicillin/streptomycin. Cells were maintained at 37 °C in 5% $CO_2$. For murine podocyte cultures, isolated glomeruli were subjected to a second enzymatic digestion with collagenase type II for 30 min at 37ºC to obtain a single-cell suspension enriched in podocyte. Cells were centrifugated, resuspended in the appropriate culture medium and maintained at 37 °C in 5% $CO_2$. Cells were subsequently trypsinized and used for downstream experiments.

*Podocyte conditioned medium and cross-talking experiment*

Conditioned medium from human or murine podocytes was collected from fully differentiated cells exposed to ADR (1µg/ml) for 24 hours. The resulting conditioned supernatant was subsequently applied to human and murine PECs for 24 hours to evaluate the molecular responses.

<u>Fetal Bovine Serum (FBS) extracellular vesicle free</u>

Once decomplemented at 55 °C for 30 minutes, FBS was aliquoted into polycarbonate centrifuge bottles (355618, Beckman Coulter). The samples were then ultracentrifuged using a 70Ti rotor (Beckman Coulter) in an ultracentrifuge model Optima XE-90 at 100,000 g for 22 hours at 4 °C. After ultracentrifugation, the supernatant was carefully collected and used to supplement the culture medium for subsequent stimulation experiments (only used in secondary senescence experiments).

**Senescence-Associated-β-Galactosidase Activity**

Cells were seeded into 24-well plates at a density of 50,000 cells/well. Murine podocytes were exposed to ADR 1µg or ADR 1µg + DQ (D;600nM, Q;30µM) for 24h, while murine PECs were treated with the podocyte conditioned medium. After 24h, the senescence-associated-β-galactosidase (SA-β-gal) activity was measured using the Senescence Cells Histochemical Staining Kit (CS0030, Sigma-Aldrich, St. Louis, MO, USA), as previously described[4]. The ratio of SA-β-gal positive cells/total cells was calculated by treatment-blinded manual counting of at least 1000 total cells in 5 photographs EVOS M5000 imaging system (Invitrogen, Waltham, MA, USA).

**Animals**

Male and female BALB/c mice (12–14 weeks old) were obtained from Charles River Laboratories (Barcelona, Spain) and housed under specific pathogen-free conditions at the IIS-Fundación Jiménez Díaz Animal Facility. Animals had free access to standard chow and water and were maintained under controlled temperature and humidity conditions with a 12-hour light/dark cycle. Mice were randomly assigned to the experimental groups. All animal procedures were performed in accordance with the European Union Directive 2010/63/EU and the ARRIVE guidelines for the care and use of laboratory animals. Experimental protocols were approved by the Animal Ethics Committee of IIS-Fundación Jiménez Díaz and the Comunidad de Madrid (PROEX 0019.0/26).

**Adriamycin-Induced FSGS Model and Senolytic Treatment**

Experimental focal segmental glomerulosclerosis (FSGS) was induced by a single retro-orbital injection of adriamycin (ADR; 10 mg/kg body weight) dissolved in 0.9% saline solution. Mice received the senolytic combination of dasatinib plus quercetin (DQ; dasatinib 5 mg/kg and quercetin 50 mg/kg body weight) or vehicle (10% polyethylene glycol; Sigma-Aldrich, St. Louis, MO, USA) by oral gavage. Treatment was initiated 3 days after ADR administration and continued every 4 days until sacrifice at the predefined endpoints (2 or 3 weeks). Administration of the vehicle alone did not induce detectable renal alterations. Animals were randomly allocated to four experimental groups: Control, DQ, ADR, and ADR+ DQ. The ADR-induced nephropathy model in BALB/c mice is a well-established experimental model that reproduces key pathological features of human FSGS[5,6]. The DQ dosing regimen was selected based on the pharmacokinetic characteristics of both compounds[7,8] and previous studies demonstrating effective senolytic activity in vivo[9,10] and studies in models of renal damage[11].

**Sample Collection and Biochemical Analyses**

Mice were euthanized with 5 mg/kg xylazine (Rompun,Bayer AG, Leverkusen, Germany) and 35 mg/kg ketamine (Ketolar, Pfizer, New York City,NY, United States). Blood samples were collected, and kidneys were perfused in situ with cold saline before harvesting. One half of each kidney was fixed in formalin, paraffin-embedded, and processed for histological and immunohistochemical analyses, whereas the remaining tissue was snap-frozen in liquid nitrogen and stored at −80°C for RNA and protein studies. Serum creatinine concentrations was measured using a commercially available colorimetric assay kit (Creatinine Assay Kit, ab65340; Abcam, Cambridge, UK) according to the manufacturer's instructions.

**Histology, Immunohistochemistry and Immunofluorescence**

Kidney tissues obtained from experimental mice and human kidney biopsies were fixed in 4% formaldehyde, paraffin-embedded, and sectioned at 3 µm thickness for mouse kidneys and at 1 µm thickness for human biopsies.  Sections were used for Periodic acid-Schiff (PAS, Sigma-Aldrich) as described elsewhere[4], immunohistochemistry (IHC), or immunofluorescence (IF) analyses.

**Immunohistochemistry (IHC)**

Tissue sections were deparaffinized and rehydrated through graded alcohols. Antigen retrieval was performed using sodium citrate buffer (10 mM, pH 6.0 or 9.0) in a PT Link system (Dako,

Glostrup, Denmark). Endogenous peroxidase activity was blocked with 3% hydrogen peroxide (Millipore, Burlington, MA, USA), and non-specific binding was prevented using casein blocking solution (Vector Laboratories, Newark, CA, USA). Sections were incubated overnight at 4°C with primary antibodies p21Cip1 (1:200, ab188224, Abcam, Cambridge, UK) and CD44 (1:200, ab119348 Abcam, Cambridge, UK), followed by incubation with biotinylated secondary antibodies (anti-rabbit or anti-rat, 1:200;) and avidin-biotin complex (ABC; Vector Laboratories, Newark, CA, USA). Immunoreactivity was visualized using 3,3′-diaminobenzidine (DAB; Abcam, Cambridge, UK) as chromogen. Slides were counterstained with Carazzi's hematoxylin, dehydrated, and mounted with DPX mounting medium (Merck, Darmstadt, Germany). Quantitative analysis was performed using Image-Pro Plus software v7.1 (Media Cybernetics, Rockville, MD, USA). Positive staining was quantified as the percentage of stained area relative to the total tissue area in 5–10 randomly selected fields per section (×200 magnification). For p21Cip1 staining, positive cells were manually counted in the same number of fields. IHC in human kidney biopsies for $p21^{Cip1}$ was performed using a Ventana RTU antibody DCS-60.2 (Cell Marque, Rocklin, CA, USA).

**Immunofluorescence (IF)**

Cells were sown in p6 well-plates in which previously cover glass standard line (Labbox) were added. Cells were then fixed with PFA 4% followed by several washes, first with glycine 100mM during 10 minutes and than with PBS1X. Cells were then treated as tissue sections. For IF, renal sections were deparaffinized, rehydrated, and subjected to antigen retrieval as described in IHC. Sections and cells were then permeabilized with 0.2% Triton X-100 in PBS and blocked with 4% normal rabbit (S-5000, Vector Laboratories), rat (S24-M, Sigma-Aldrich) or goat (S-1000, Vector Laboratories) serum, as appropriate, diluted in 4% bovine serum albumin (BSA, 9048-46-8, Thermo Scientific) in PBS. Cells and tissue sections were incubated overnight at 4ºC with primary antibodies diluted in TBS with 2% serum and 4% BSA: anti-p21cip1 (1:200, ab188224, Abcam, Cambridge, UK), rat anti-Nephrin (1:200 PA5-20330, Invitrogen), anti-CD44 (1:200, ab119348 Abcam, Cambridge, UK) and phalloidin (1:1000, SC-363794, Santa Cruz; CA, USA). Samples were incubated with the secondary antibodies included Alexa Fluor 488-conjugated anti-rabbit IgG (1:250; S-5000 Thermo Fisher Scientific, Waltham, MA, USA) and Alexa Fluor 563-conjugated anti-rat IgG (1:250, A21208; Thermo Fisher Scientific, Waltham, MA, USA).

Nuclei were counterstained with 4',6-diamidino-2-phenylindole (DAPI; 1:10,000; Sigma-Aldrich, St. Louis, MO, USA). Sections were mounted using ProLong Gold Antifade Reagent (Invitrogen, Waltham, MA, USA) and images were acquired at 40x with a BX53 fluorescence microscope (Olympus, Tokyo, Japan).

**Glomerular injury score**

The severity of focal segmental glomerulosclerosis in mouse kidneys was assessed semiquantitatively in PAS-stained sections by an expert renal pathologist, blinded to group allocation. All glomeruli present in each section were evaluated by visual inspection, and an overall severity score was assigned per section as follows: 0 (no abnormalities/within normal limits), 1 (mild sclerosis in at least some glomeruli), 2 (moderate sclerosis in at least some glomeruli), or 3 (severe sclerosis in at least some glomeruli).

**Protein level studies**

Total protein from frozen renal cortex and pelleted cultured cells were isolated by homogenization in a lysis buffer (50 mM of Tris-HCl, 150 mM of NaCl, 2 mM of EDTA, 2 mM of EGTA, 0.2 % Triton X-100, 0.3 % NP-40)) with 10 µL/mL protease inhibitor cocktail (P8340, Sigma-Aldrich) , 0.2 mmol/L phenylmethylsulfonyl fluoride (PMSF, P7626, Sigma-Aldrich), and 0.2 mmol/L sodium orthovanadate (S-6508, Sigma-Aldrich). Proteins were quantified using a Pierce BCA protein assay kit (Thermo Fisher Scientific; Waltham, MA, USA) and then were separated by electrophoresis using 8–10% polyacrylamide-SDS gels under reducing conditions for western blotting. Samples were then transferred onto polyvinylidene difluoride membranes (Thermo Fisher Scientific; Waltham, MA, USA), blocked with 5% non-fat milk, and incubated overnight at 4°C with primary antibodies : anti-CDKN1A/p21 (1:200, sc-6246 Santa Cruz; CA, USA), anti-phospho(S32)-IκBα (1:1000; sc-8404, Santa Cruz; CA, USA), anti-pERK (1:200; sc-7383 Santa Cruz; CA, USA), anti-ERK (1:1000; #4370, Cell Signaling; MA, USA), anti- pSRC (1:1000; #6943, Cell Signaling; MA, USA), anti-SRC (1:1000; #2109, Cell Signaling; MA, USA), anti-p62 (1:1000; #5114S, Cell Signaling; MA, USA), anti-pSTAT3 (1:1000; #9134, Cell Signaling; MA, USA), anti STAT3 (1:1000; #8768, Cell Signaling; MA, USA), anti-LC3B (1:200;NB100-220, Novus Biologicals) and,anti-GAPDH antibody (1:5000; CB1001, Millipore; MA, USA). Then, the membranes were incubated with HRP-conjugated secondary antibodies anti-rabbit (1:5000, NA934VS, Amersham) or anti-mouse (1:5000, NA931VS, Amersham). Loading controls were performed using a mouse anti-GAPDH antibody (1:5000; CB1001, Millipore; MA, USA). Proteins on membranes were visualized using the chemiluminescence detection kit Immobilon Crescendo Western HRP substrate (Millipore; Burlington, MA, USA) on the Amersham Imager 600 instrument (GE Healthcare; Chicago, IL, USA). Images were analyzed by densitometry using the image processing program written on Java, ImageJ.

**Proteomics analysis by liquid chromatography coupled to mass spectrometry**
**Sample preparation**

Renal tissues from the ADR (n = 4), ADR+ D&Q (n = 5) and control (n = 4) groups were processed for proteomic analysis. Renal cortex tissues (9-18 mg) were homogenized with a lysis buffer containing 2.5% sodium dodecyl sulfate (SDS), 50 mM triethylammonium bicarbonate (TEAB), 5 mM tris(2-carboxyethyl) phosphine (TCEP), and 10 mM iodoacetamide (IAA). Kidney lysates were treated with DNase (2500 U/mL; Thermo Scientific™, Cat. No. 90083) at 37 °C for 15 minutes to remove contaminating DNA and incubated at 60 °C for 30 minutes to reduce and alkylate cysteine disulfide bonds. Samples were sonicated using a UP50H ultrasonic processor (Hielscher Ultrasonics) for 1 minute on ice (0.5 cycles, 100% amplitude), followed by centrifugation at 18,400 × g for 10 min. Total protein was measured using the Pierce 660 nm colorimetric assay (Thermo Scientific) supplemented with ionic Detergent Compatibility Reagent (Thermo Scientific). Protein digestion was performed using a solid-phase-enhanced sample preparation (SP3)-based workflow with MagReSyn® Hydroxyl beads (Resyn Biosciences), automated on an Opentrons OT-2 liquid-handling robot, according to a previously established protocol[12]. Briefly, 60 µg of each sample was automatically processed on the Opentrons OT-2 platform. Protein digestion was performed overnight at 37 °C using trypsin and Lys-C at enzyme-to-protein ratios of 1:33 and 1:500, respectively, in a Jitterbug 4-plate incubated microplate shaker (Boekel Scientific). The eluted peptides were dried in a speed vacuum and quantified by fluorimetry (QuBit) according to the manufacturer's instructions.

**Analysis by liquid chromatography coupled to mass spectrometry OE-240**

For nanoliquid chromatography coupled with Electrospray Ionization Tandem Mass Spectrometry (nanoLC-ESI-MS/MS) analysis, 500 ng of each sample was individually analyzed using an Ultimate 3000 nano HPLC system (Thermo Fisher Scientific) coupled online to an Orbitrap Exploris™ 240 mass spectrometer (Thermo Fisher Scientific). Each sample (500 ng in 5 µL of sample resuspended in mobile phase A: 0.1% formic acid (FA)) was loaded on a 50 cm × 75 µm µPAC Neo C18 analytical column at 50°C and separated at a flow rate of 250 nL/min using a 120 min gradient ranging from 2% to 95% mobile phase B (80% acetonitrile (ACN) in 0.1% FA). In addition, an internal standard consisting of a pool of all replicates for each condition was run between sample batches. To avoid carryover, two 60-min blank samples (mobile phase A) were systematically run between samples. Data acquisition was performed in data-independent acquisition mode with variable mass-to-charge ratio (m/z) windows.

**Proteomics Data Analysis**

Raw DIA data files were analyzed using Spectronaut (version 20.2, Biognosys AG, Schlieren, Switzerland) in DirectDIA+ (Deep) mode, which integrates complex DIA spectra and performs database searching via the Pulsar search engine. The search was conducted against a *Mus musculus* UniProtKB database, which contains common laboratory contaminants. The Pulsar search engine generated an in silico spectral library from DIA data for peptide and protein identification. The search mode was set to Closed Search, with the following parameters: cysteine carbamidomethylation (+57.021464 Da) as fixed modification and methionine oxidation (M) (+15.994915 Da), N-term acetylation (+42.010565 Da), and Gln→pyro-Glu (-17.026549 Da) as variable modifications. Tolerance was applied according to DirectDIA+ (Deep) workflow default parameters, and Trypsin/P was selected as a protease, with a maximum of two missed cleavages. To ensure high identification confidence, false discovery rate (FDR) was applied at the PSM, peptide, and protein levels (FDR<1%). Quantitative information was extracted from MS2 signals according to area to estimate relative peptide and protein abundances across samples. The precursor filtering mode was set to Identified (q-value). Synchronized retention time alignment was performed to ensure consistent peptide quantification across runs. Background noise removal was enabled, and group q-value filtering was used for multi-channel datasets. Intensity values were exported and normalized in R using the limma package[13] applying the Cyclic-Loess method to correct for systematic biases between samples. Missing values were subsequently imputed using a hybrid strategy based on the presumed missingness mechanism, combining minimum-value replacement (Min) with robust sequential imputation using ImpSeqRob (MAR). A linear modeling framework was then fitted for differential analysis, and the empirical Bayes (eBayes) approach was used to moderate standard errors. Proteins with adjusted p-values ≤ 0.05 and corresponding $\log_2$ fold-change ($\log_2$FC) values were considered differentially abundant. Functional enrichment analysis was performed using Gene Set Enrichment Analysis (GSEA) v4.4.0 on a pre-ranked gene list derived from the differential expression results[14–16]. Functional enrichment analysis of differentially abundant proteins (*p-value* < 0.05) was performed using Metascape, based on Gene Ontology (GO), KEGG, and Reactome pathway annotations[17].

**Gene Expression assays**

Total RNA was isolated from frozen renal cortex and pelleted cultured cells by homogenization with TRItidy G (PanReac; Darmstadt, Germany), according to the manufacturer's protocols. Next, cDNA was synthesized using a High-Capacity cDNA Reverse Transcription Kit (Applied Biosystems; Waltham, MA, USA) using 2 µg of total RNA. Gene expression analysis was determined by quantitative PCR (qPCR) using the commercial master mix Premix Ex Taq (Takara; Otsu, Japan) and predesigned TaqMan-based qPCR assays for target and housekeeping genes (Table 1) and run on QuantStudio 3 thermocycler (Applied Biosystems; Waltham, CA, USA). Relative expression values were normalized by the expression of housekeeping genes using the $2^{-\Delta\Delta Ct}$ method. The results were expressed as fold change (n-fold) relative to control. In the heatmaps, row Z-scores were calculated for each gene by standardizing its relative expression values across samples.

**Table 1. Pre-designed assays used for qPCR.**

| Specie | Gene | Gene type | Assay ID |
|---|---|---|---|
| Mouse | *Cdkn1a* | Target | Mm00432448_m |
| Mouse | *Cdkn2a* | Target | Mm00494449_m1 |
| Mouse | *Ccl2* | Target | Mm00441242_m1 |
| Mouse | *Ccl5* | Target | Mm01302428_m1 |
| Mouse | *Ccl8* | Target | Mm01297183_m1 |
| Mouse | *Cd44* | Target | Mm01277161_m1 |
| Mouse | *Cd9* | Target | Mm00514275_g1 |
| Mouse | *Mki67* | Target | Mm01278617_m1 |
| Mouse | *Il6* | Target | Mm00446190_m1 |
| Mouse | *tgf-β1* | Target | Mm00441726_m1 |
| Mouse | *Cxcl12* | Target | Mm00445553_m1 |
| Mouse | *Atg5* | Target | Mm01187303_m1 |
| Mouse | *Atg7* | Target | Mm00512209_m1 |
| Mouse | *Becn1* | Target | Mm01265461_m1 |
| Mouse | *Bcl2l1l* | Target | Mm00437783_m1 |
| Mouse | *Csf1* | Target | Mm00432691_m1 |
| Mouse | *B2m* | Target | Mm00437762_m1 |
| Mouse | *Ifng* | Target | Mm01168134_m1 |
| Mouse | *Ifit1* | Target | Mm00515153_m1 |
| Mouse | *Mx1* | Target | Mm00487796_m1 |
| Mouse | *Isg15* | Target | Mm01705338_s1 |
| Mouse | *Oasl2* | Target | Mm01201449_m1 |
| Mouse | *Ifna* | Target | Mm02525960_s1 |
| Mouse | *Gapdh* | Housekeeping | Mm99999915_g1 |
| Human | *CCL2* | Target | Hs00234140_m1 |
| Human | *CCL5* | Target | Hs00982282_m1 |
| Human | *TGF-β* | Target | Hs00998133_m1 |
| Human | *CDKN1A* | Target | Hs00355782_m1 |
| Human | *CDKN2A* | Target | Hs00923894_m1 |
| Human | CSF-1R | Target | Hs00911250_m1 |
| Human | *CSF-1* | Target | Hs00174164_m1 |
| Human | *CCN2* | Target | Hs00170014_m1 |
| Human | *CXCL12* | Target | Hs03676656_mH |
| Human | *CD44* | Target | Hs00153304_m1 |
| Human | *IL6* | Target | Hs00174131_m1 |
| Human | *CLDN1* | Target | Hs00221623_m1 |
| Human | *IFI44C* | Target | Hs00197427_m1 |
| Human | *IFIT1* | Target | Hs03027069_s1 |
| Human | *IFI27* | Target | Hs00271467_m1 |
| Human | *ISG15* | Target | Hs01921425_s1 |
| Human | *GAPDH* | Housekeeping | Hs02786624_g1 |

**Statistical Analysis.**

Kidneys from all groups were compared to control, expressing results as fold-change over control values of 1. Results are expressed as fold increase with respect to the control average as mean ± standard error of the mean (s.e.m) of 3 to 13 animals per group.

The Shapiro–Wilk test was used to evaluate sample normality distribution. To assess statistical differences between groups, several tests were used based on data characteristics. When samples followed the Gaussian distribution, a one-way ANOVA followed by the Fisher's LSD test was used. To compare non-parametric samples, a Kruskal–Wallis and a subsequent Uncorrected Dunn's test analysis were performed.

For isolated glomeruli experiments t-test was done. For cell culture experiments, paired comparisons were done using RM one-way ANOVA, with the Geisser-Greeehouse correction for normal data or Friedman test for non-normal data, followed by Fisher's LSD or Dunn's test, respectively. All statistical analyses and graphs were performed using GraphPad Prism 9.0 (GraphPad Software, San Diego, CA, USA). *P*-values < 0.05 were considered statistically significant.
